# Tuft cells mediate commensal remodeling of the small intestinal antimicrobial landscape

**DOI:** 10.1101/2022.10.24.512770

**Authors:** Connie Fung, Lisa M. Fraser, Gabriel M. Barrón, Matthew B. Gologorsky, Samantha N. Atkinson, Elias R. Gerrick, Michael Hayward, Jennifer Ziegelbauer, Jessica A. Li, Katherine F. Nico, Miles D.W. Tyner, Leila B. DeSchepper, Amy Pan, Nita H. Salzman, Michael R. Howitt

## Abstract

Succinate produced by the commensal protist *Tritrichomonas musculis* (*T. mu*) stimulates chemosensory tuft cells, resulting in intestinal type 2 immunity. Tuft cells express the succinate receptor SUCNR1, yet this receptor does not mediate anti-helminth immunity nor alter protist colonization. Here, we report that microbial-derived succinate increases Paneth cell numbers and profoundly alters the antimicrobial peptide (AMP) landscape in the small intestine. Succinate was sufficient to drive this epithelial remodeling, but not in mice lacking tuft cell chemosensory components required to detect this metabolite. Tuft cells respond to succinate by stimulating type 2 immunity, leading to interleukin-13-mediated epithelial and AMP expression changes. Moreover, type 2 immunity decreases the total number of mucosa-associated bacteria and alters the small intestinal microbiota composition. These findings demonstrate that a single metabolite produced by commensals, like *T. mu*, can markedly shift the intestinal AMP profile and suggest that tuft cells utilize SUCNR1 to modulate bacterial homeostasis.

## Introduction

The small intestine (SI) provides a massive surface area for the digestion and absorption of ingested nutrients. At the same time, it contends with continuous interactions with numerous pathogenic and commensal microbes, where endogenous bacterial densities can reach as high as 10^7^ - 10^8^ cells per gram of contents in the distal SI, or ileum (1, 2). A mucus layer permeated with antimicrobial peptides (AMPs) forms a chemical barrier that prevents these microbes from overgrowing, contacting the epithelium, and/or translocating across mucosal tissues (3–5). Loss of this barrier results in excessive inflammatory responses to the microbiota and is a pathophysiological feature of many gastrointestinal diseases (6).

While mucus and AMPs influence gut microbiota composition and localization (5, 7–9), microbes reciprocally play a critical role in forming and maintaining proper barrier function within the intestinal tract. Germ-free animals have fewer mucus-producing goblet cells, a thinner mucus layer, and reduced production of specific AMPs (10, 11). Strikingly, colonization of germ-free mice with a conventional microbiome is sufficient to restore conventional mucus properties and Paneth cell expression of specific AMPs within these animals (10, 12–14). This induction of specific AMP production by the microbiota occurs via pattern-recognition receptor signaling (15) or through sensing microbially-derived metabolites, such as short-chain fatty acids (16).

Much of our knowledge about how the gut microbiota modulates barrier function and intestinal physiology derives from studies on commensal bacteria. However, this microbial ecosystem also includes archaea, fungi, viruses, and protists (17, 18). Recently, commensal protists from the genus *Tritrichomonas* have emerged as important members of the gut microbiota of wild and laboratory rodents (19). One of these protist species, *Tritrichomonas musculis* (*T. mu*), induces profound changes to the murine gut epithelium and immune system. Type 2 immunity, the hallmark immune response mounted against parasitic worm infections, is activated in the distal SI in response to *T. mu*, concomitant with a notable expansion of specialized chemosensory epithelial cells, called tuft cells, and mucus-producing goblet cells (20). Succinate, a metabolic byproduct of commensal tritrichomonads, activates tuft cell taste-chemosensory signal transduction by engaging the G-protein-coupled receptor SUCNR1 (21– 23). In response to succinate, tuft cells release the cytokine interleukin-25 (IL-25), which activates group 2 innate lymphoid cells (ILC2) in the lamina propria. Subsequently, ILC2s produce the type 2 cytokine interleukin-13 (IL-13), which initiates type 2 immune responses within the gut. In addition, IL-13 biases intestinal stem cells to differentiate into tuft or goblet cells, thereby amplifying this tuft cell-ILC2 circuit (20, 24, 25).

Succinate is an intriguing tuft cell-activating ligand, as many organisms, including commensal bacteria, tritrichomonads, parasitic worms, and mammalian host cells produce this metabolite (22, 23, 26–28). Thus, luminal accumulation of succinate does not distinguish between host or microbe, nor commensal or pathogen. In fact, the succinate receptor SUCNR1 is dispensable for detecting helminths and initiating anti-parasitic immunity (21, 22). Furthermore, intestinal type 2 immunity triggered by *T. mu*-produced succinate does not reduce or enhance protist colonization (20, 22). These observations suggest that tuft cell detection of succinate and the associated changes to the gut epithelial and immunological landscape has not evolved for anti-parasitic responses. Interestingly, SUCNR1 expression is particularly enriched on SI tuft cells compared to other parts of the gastrointestinal tract (22, 29). Furthermore, succinate induces the most pronounced tuft cell hyperplasia in the distal SI (ileum) compared to the proximal SI (22), suggesting that ileal tuft cells are particularly sensitive to luminal succinate. However, why ileal tuft cells orchestrate type 2 immunity in response to this metabolite is still not well understood.

To address this question, we profiled the global effects of microbial-derived succinate on ileal physiology. We used spatial transcriptomics (ST) as an unbiased approach to characterize the impact of *T. mu*, a major succinate producer, on distal SI gene expression *in situ. T. mu* induced substantial transcriptional changes in the SI epithelium and lamina propria, including increased goblet and Paneth cell numbers and profound alterations to the antimicrobial expression profile. Furthermore, we demonstrated that these shifts in AMP production are dependent on tuft cell detection of *T. mu*-produced succinate and subsequent activation of type 2 immunity. Finally, we discovered that type 2 immune induction significantly decreases the total number of mucosa-associated bacteria in the distal SI, most notably segmented filamentous bacteria (SFB). This suggests increased microbial killing and restructuring of the gut community by upregulated AMPs. Collectively, our work highlights the significant impact succinate exerts on the cellular composition of the gut epithelium and host-microbiota interactions through activating tuft cells.

## Results

### The commensal protist *T.mu* induces significant remodeling of the intestinal epithelium

To map the impact of microbial-derived succinate on the SI landscape, we conducted spatial transcriptomics (ST) on uncolonized and *T. mu*-colonized mice. We first confirmed that although *T. mu* colonization slightly increases tuft cell numbers in the proximal SI, the increase is much more pronounced in the distal SI (Fig. S1A), consistent with prior studies showing similar patterns with succinate (22). Consequently, we focused on the ileum for our ST analysis. We harvested ileal tissue from one uncolonized mouse and one mouse colonized with *T. mu* for 3 weeks and affixed 10 μm sections to Visium slides (10x Genomics). These slides have thousands of 55 μm-diameter spots that each contain mRNA-binding probes with a unique barcode. A single spot captures the gene expression of approximately 5-15 cells within the tissue section (Fig. 1A). This approach produced complete transcriptomes of the cell populations within each ST spot and allowed us to map these gene expression signatures *in situ* (Fig. S1B).

**Figure 1.**
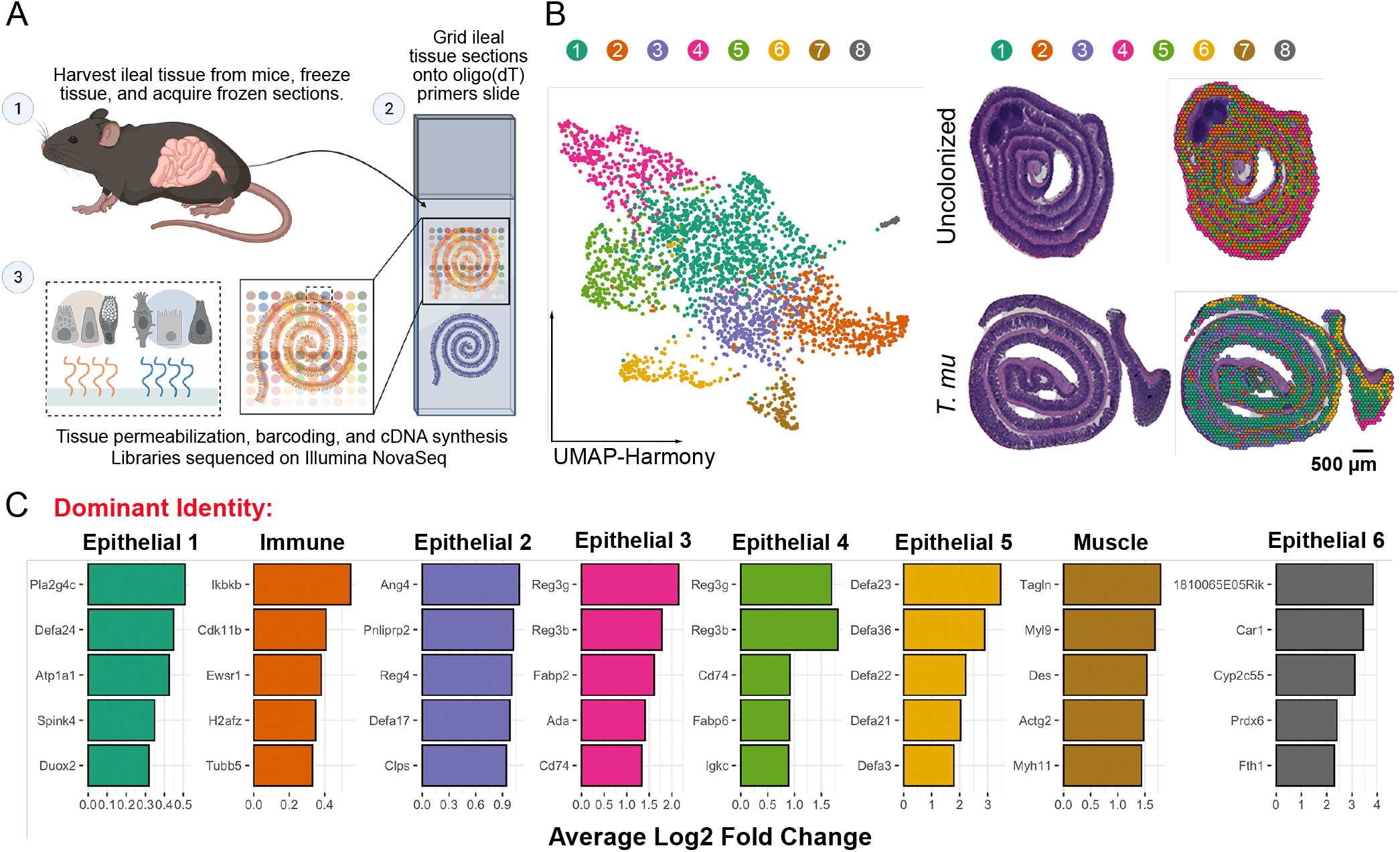
Spatial transcriptomics (ST) reveals that *T. mu* colonization induces profound transcriptional changes to the small intestinal epithelium. (A) Schematic of ST pipeline. (B) Harmonized UMAP plots of ST spots organized by cluster identity (left); corresponding hematoxylin and eosin (H&E)-stained tissue scans with overlaid Seurat clustering (right). (C) Top 5 highly expressed genes for each cluster shown by log 2-fold change compared to other clusters. Clusters annotated based on dominance of gene expression associated with epithelial, immune, or muscle cells.

Using the ST analysis tool, STUtility (30), we manually annotated the tissue-associated spots with one of three identities: (1) Muscularis (Muscle), (2) Peyer’s Patch (PP), or (3) Epithelium/Lamina Propria (Epi/LP) (Fig. S1C). We excluded the muscularis and Peyer’s patch compartments for downstream analysis to focus on epithelial and immune changes induced by *T. mu*. Taking the Epi/LP spots, we performed non-negative matrix factorization (NNMF) to create gene and cell topic modules that described the two samples (Fig. S1D). We then used these factors to integrate the sequencing data from the two samples via Harmony (31) and represent this as eight distinct clusters, as visualized by UMAP projection (Fig. 1B). We also mapped these clusters back onto the tissue array to visualize their spatial location within the ileum (Fig. 1B). Since each dot on the UMAP projection represents a spot comprised of multiple cells, we could not assign a specific cellular identity to these clusters. Therefore, we annotated the clusters based on their dominant expression of genes associated with epithelial, immune, or muscle function (Fig. 1C). *T. mu* colonization highly altered the abundance and composition of all clusters, except for Cluster 6, which is enriched with muscle and stromal genes (i.e. *Tagln, Myl9 and Myh11*) (Fig. 1B-C and S1E). Because *T. mu* is known to induce tuft cell hyperplasia in the ileum (20, 22, 23), we confirmed an increase in spots containing the tuft cell-specific genes *Trpm5* and *Dclk1* in the *T. mu*-colonized sample relative to the uncolonized sample (Fig. S1F). We also stained serial sections with anti-DCLK1 antibodies and found increased tuft cell abundance in the *T. mu*-colonized ileum by microscopy (Fig. S1G).

### Integration of single-cell RNA sequencing (scRNA-seq) with ST identifies hybrid goblet-Paneth cell signatures during *T. mu* colonization

Six of the seven clusters altered by *T. mu* were epithelial-dominant (Fig. 1C). To identify the epithelial composition of each ST spot within our samples, we anchored a previously published reference scRNA-seq dataset of the SI epithelium onto our spatial analysis (32). This approach allowed us to deconvolute the transcriptomes of the ST spots with the 15 annotated epithelial cell types present in the reference scRNA-seq dataset (Fig. 2A). We calculated the probability of a specific epithelial cell identity within each spot; non-epithelial cell identities were excluded from this analysis. To visualize the deconvoluted spots in a spatial context, we represented the proportion of cell types per spot as pie charts arranged on the tissue array (Fig. 2B-C). Immature and mature distal enterocyte identities predominated the spots in the uncolonized sample, whereas goblet and Paneth cell identities were the most prevalent in the *T. mu*-colonized sample (Fig. S2A-C). The increased goblet cell expression signatures was expected, as goblet cell hyperplasia was previously reported with tritrichomonad colonization (20, 22, 23). However, an increase in Paneth cell expression signatures in response to *T. mu* has not been described.

**Figure 2.**
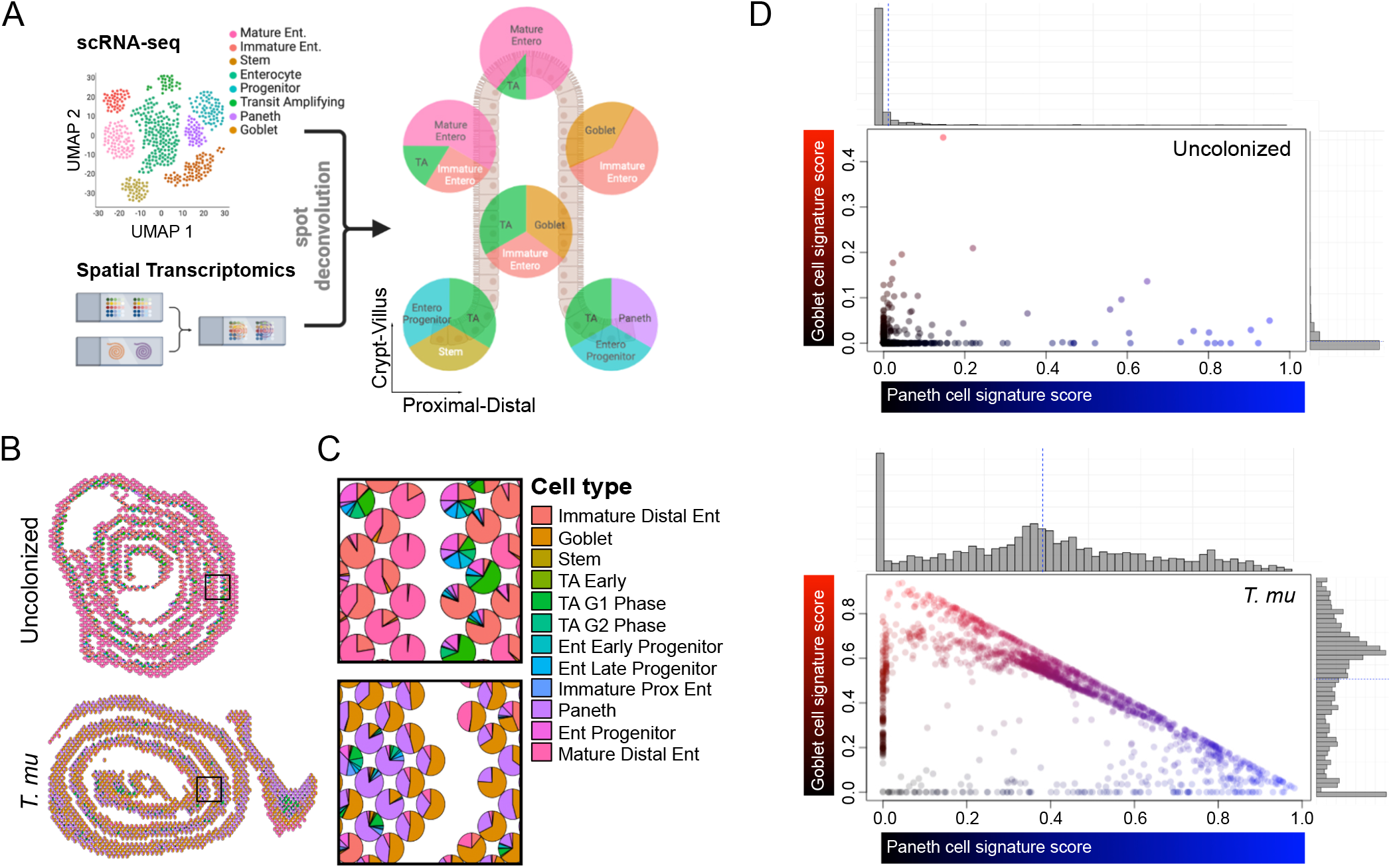
Single-cell RNA sequencing (scRNA-seq) integration with ST reveals goblet and Paneth cell expansion in the *T. mu*-colonized small intestine. (A) Schematic of Seurat anchoring method to integrate a single-cell RNA sequencing (scRNA-seq) small intestinal epithelial dataset (32) with the ST dataset. (B and C) Scatter pie plots of the ileum depicting predicted cellular identities (B), with corresponding zoomed regions of interest (C). Ent, Enterocyte; TA, Transit-amplifying; G1, G1/S cell-cycle phase; G2, G2/M cell-cycle phase; Prox, Proximal. (D) Visualization of goblet (red) and Paneth cell (blue) signature scores of ST spots colored by identity. Gray histograms on top and right show distribution of spots along each axis.

Paneth cells are typically restricted to the intestinal crypts, while goblet cells are distributed throughout the crypt-villus axis (33, 34). Nonetheless, because Paneth cell and goblet cell expression signatures were so highly enriched in the *T. mu*-colonized sample (Fig. 2C and S2A-C), we investigated whether the two cell types collocate in the distal SI and if they share the same cellular neighborhoods. We plotted the spots in a two-dimensional space, where the axes represent the percentage of Paneth or goblet cell signature of each ST spot. Most spots from the uncolonized ileum aligned along either the x- or y-axis, indicating a binary goblet or Paneth cell identity. In contrast, spots from the *T. mu*-colonized ileum were distributed across the spectrum of shared identities between goblet and Paneth cells (Fig. 2D). In addition, these hybrid goblet-Paneth cell spots localized throughout the crypt-villus axes (Fig. S2A). This indicates that *T. mu* colonization may: 1) increase the number of Paneth cells residing outside of the crypts among villus goblet cells, or 2) increase the abundance of a cellular subtype that expresses both Paneth and goblet cell features. While our ST data does not distinguish between either possibility, we observed cells above the crypts that are reminiscent of “intermediate cells” in *T. mu*-colonized tissue (Fig. S3A-B). Intermediate cells were described in the intestines as early as the 1890s and are typically rare within the gut epithelium under homeostatic conditions. They are characterized by the presence of granules within mucus globules, morphological features that are both Paneth and goblet cell-like (Fig. S3A-B) (35). Notably, helminth infections, which also activate type 2 immune responses, can induce Paneth cell hyperplasia and increase the number of “intermediate cells” in the SI (36–38). Whether these intermediate cells represent a transitionary state between Paneth and goblet cells or if they form a stable epithelial subset with specialized functions during type 2 immunity remains unclear. Altogether, these results demonstrate that *T. mu* colonization favors a secretory epithelial cell program, including a significant increase in goblet and Paneth cell gene expression signatures. *T. mu* colonization may also increase the abundance of an “intermediate cell” that morphologically resembles goblet and Paneth cells.

### *T. mu* colonization broadly alters the antimicrobial landscape in the small intestine

To understand how *T. mu* colonization remodels the epithelial and immune compartments in the distal SI, we performed differential gene expression analysis between the Epi/LP-associated clusters. We found that some of the most differentially regulated genes between the uncolonized and *T. mu*-colonized ilea encode AMPs. Surprisingly, not all AMPs are globally upregulated in the *T. mu*-colonized sample despite an increase in goblet and Paneth cell signatures in the tissue. A select group of AMPs was downregulated by *T. mu* colonization (Fig. 3A-C), with the most dramatic effects on *Reg3b* and *Reg3g*. These two AMPs belong to the Regenerating islet-derived (Reg) family of proteins, (39), are expressed by intestinal epithelial cells, and play important roles in host-microbe interactions (5, 40, 41). Innate recognition of bacteria or microbe-associated molecular patterns induce *Reg3b* and *Reg3g* expression (12, 15, 41, 42), but little is understood about stimuli that downregulate the expression of these AMPs.

**Figure 3.**
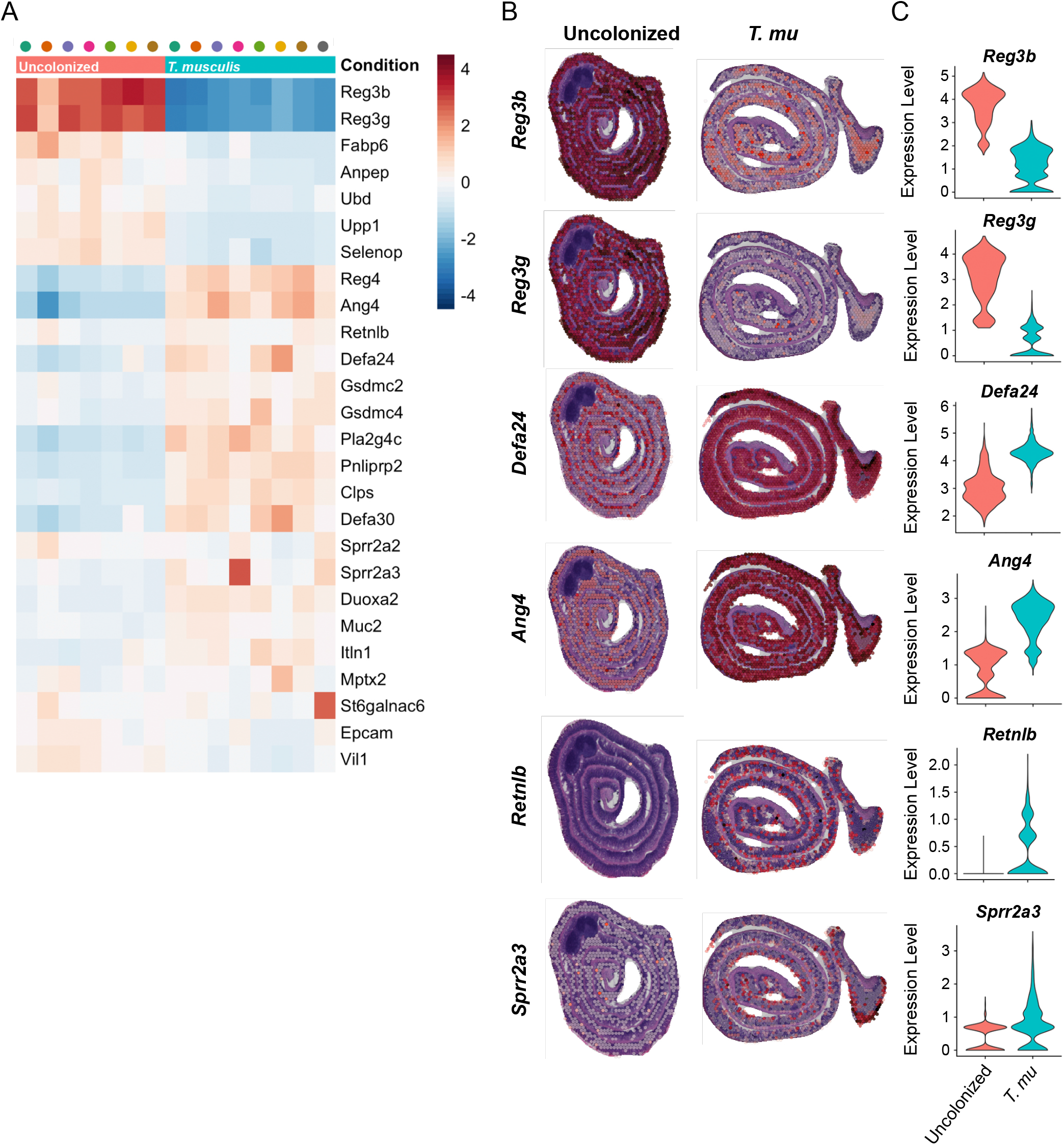
*T. mu* colonization alters the small intestinal AMP repertoire. (A) Heatmap of AMP genes clustered by sample condition and ranked with corresponding Z-score. Each column represents a cluster from the uncolonized (pink) or *T. mu*-colonized mouse (aqua). (B and C) Gene features overlay plots on ST spots throughout ileal tissue of the uncolonized or *T. mu*-colonized mouse (B), with corresponding violin plots depicting global gene expression (C).

Numerous AMP-encoding genes were also highly upregulated in response to *T. mu* colonization, including various α-defensin-encoding genes, *Ang4, Sprr2a3*, and *Retnlb* (Fig. 3A-C). The α-defensin family of AMPs are specifically produced by Paneth cells in the SI epithelium and have broad-spectrum antimicrobial activity (43). *Ang4* is a member of the RNase superfamily and is specifically produced by Paneth cells in the SI epithelium. ANG4 exhibits antimicrobial activity against specific Gram-positive and Gram-negative bacteria and fungi, and expression of this AMP is induced by the intestinal microbiota (14). *Sprr2a3* and *Retnlb* are AMPs that are upregulated by type 2 immunity. *Sprr2a3* is produced by both Paneth and goblet cells, kills Gram-positive bacteria, and is deployed during helminth infection (44). Alternatively, *Retnlb* is only expressed by goblet cells (45, 46). Resistin-like molecule β (RELMβ), the protein encoded by *Retnlb*, is induced in response to the microbiota and during helminth infections. It has both anti-bacterial and anti-helminthic properties and is critical for worm clearance (8, 45, 47). Taken together, these data illustrate that *T. mu* profoundly alters the epithelial and immune compartments of the distal SI, with broad shifts in transcriptional AMP programs. These transcriptional changes are consistent with the increases in goblet and Paneth cell signatures in *T. mu*-colonized tissue, as these two cell types are known to produce AMPs in the gut (34).

### *T. mu* alters antimicrobial production in the small intestine by activating tuft cell taste-chemosensory pathways

To investigate whether *T. mu* alters the abundance and morphology of secretory epithelial cells, we colonized WT mice with the protist for 3 weeks (Fig. S4A) and examined the distal SI by microscopy. In line with previous observations (20, 22, 23), *T. mu* colonization resulted in ileal tuft and goblet cell hyperplasia (Fig. S4B-C). Protist colonization also induced Paneth cell hyperplasia, which was associated with ultrastructural changes to these cells (Fig. 4A-C). Normal Paneth cells, as seen in uncolonized mice, contained large, electron-dense secretory granules concentrated at the apical end of the cell, with each granule surrounded by a thin, electron-lucent halo (Fig. 4C and S4D). These halos were previously shown to contain MUC2 mucins (48). Paneth cells in *T. mu*-colonized mice, however, possess features that are intermediate between Paneth and goblet cells—smaller, electron-dense granules surrounded by larger, electron-lucent mucin globules (Fig. 4C and S4D). We also visualized lysozyme, an AMP specifically produced by Paneth cells, in the ilea of uncolonized or *T. mu*-colonized mice by microscopy (33). In uncolonized mice, we found that lysozyme localized within distinct secretory granules at the apical ends of Paneth cells (Fig. 4D and S4E). However, lysozyme signal within the Paneth cells of *T. mu*-colonized mice was dimmer and dispersed throughout the entirety of the cell (Fig. 4D and S4E). Collectively, these results show that *T. mu* induces Paneth cell hyperplasia, morphological changes to Paneth cell granules, and abnormal AMP localization.

**Figure 4.**
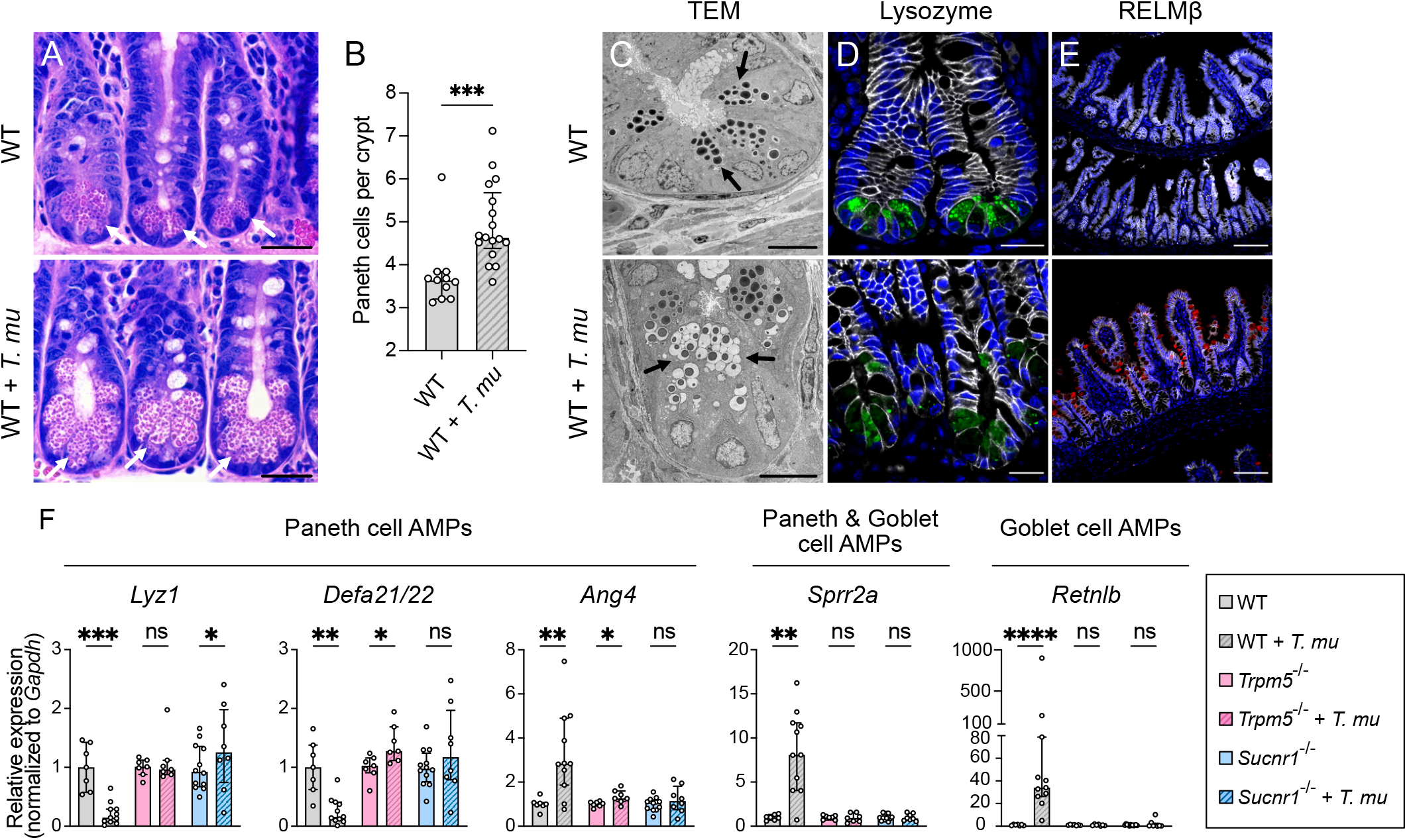
*T. mu* induces goblet and Paneth cell hyperplasia and alters AMP production in the distal small intestine via tuft cell stimulation. (A) Representative images of H&E-stained sections of the ileal crypts from uncolonized or *T. mu*-colonized WT mice. White arrows indicate representative Paneth cells. Scale bar: 25 μm. (B) Average number of Paneth cells per crypt in the ilea of uncolonized and *T. mu*-colonized WT mice (n = 11-17 mice per group). (C) Representative transmission electron microscopy images of the ileal crypts from uncolonized or *T. mu*-colonized WT mice. Black arrows indicate representative Paneth cell secretory granules. Scale bar: 10 μm. (D and E) Representative fluorescence microscopy images of the ilea from uncolonized or *T. mu*-colonized WT mice. Nuclei (blue), E-cadherin (white), Lysozyme (Paneth cell AMP) (green), RELMβ (goblet cell AMP) (red). (D) Scale bar: 20 μm. (E) Scale bar: 100 μm. (F) Expression of representative AMP genes determined by RT-qPCR in the ileal epithelial fraction from uncolonized or *T. mu*-colonized WT, *Trpm5*^-/-^, and *Sucnr1*^-/-^ mice (n = 7-12 mice per group). Relative expression normalized to *Gapdh*. In Panels B and F, center values = median; error bars = interquartile range (IQR). Significance determined using Mann-Whitney *U* tests. ns = no significance, * *p* < 0.05, ** *p* < 0.01, *** *p* < 0.001, **** *p* < 0.0001.

To determine whether *T. mu*-induced changes to AMP production occurred through tuft cell sensing of the protist metabolite succinate and subsequent activation of the taste-chemosensory pathway (20, 22, 23), we colonized WT mice, and mice either lacking TRPM5, a calcium-gated ion channel that is critical for the taste pathway (20, 49), or SUCNR1, a G-protein coupled receptor that detects succinate (22, 23, 50) (Fig. S4A). We measured the relative expression of a panel of representative AMP genes produced by Paneth and/or goblet cells via RT-qPCR. *T. mu* colonization altered AMP gene expression in the ileal epithelium, with decreased expression of *Lyz1* and *Defa21/22* (Paneth cells), and increased expression of *Ang4* (Paneth cells), *Sprr2a* (Paneth and goblet cells), and *Retnlb* (goblet cells) (Fig. 4F). These transcriptional changes are consistent with ileal lysozyme and RELMβ protein levels (encoded by *Retnlb*) in *T. mu*-colonized mice (Fig. 4E and S4F-G). In contrast to WT mice, these AMP gene expression changes were abrogated in *T. mu*-colonized *Trpm5*^-/-^ and *Sucnr1*^-/-^ mice compared to uncolonized controls (Fig. 4F and S4F). Altogether, these results suggest that *T. mu* alters AMP production in the distal SI by stimulating tuft cell taste-chemosensory pathways via the metabolite succinate.

### Succinate drives changes to intestinal AMP production by stimulating tuft cells and activating type 2 immunity

Succinate is sufficient to induce tuft and goblet cell hyperplasia in the distal SI (22, 23); however, little is known about its effect on Paneth cells. To determine if succinate is sufficient to drive changes to Paneth cell numbers, morphology, and AMP production in the ileum, we treated germ-free (GF) mice and conventional (CV) mice with 100 mM succinate in their drinking water for one week, after which tissues were collected for microscopy, RT-qPCR, and western blot analysis. Succinate increased both tuft and goblet cell numbers in GF and CV mice and led to a striking increase in MUC2 throughout the crypt-villus axes of GF and CV mice (Fig. S5A-C). In addition, we found that succinate treatment significantly increased Paneth cell numbers in both GF and CV mice compared to respective controls (Fig. 5A-B), indicating that this metabolite alone is sufficient to drive Paneth cell hyperplasia in the distal SI.

**Figure 5.**
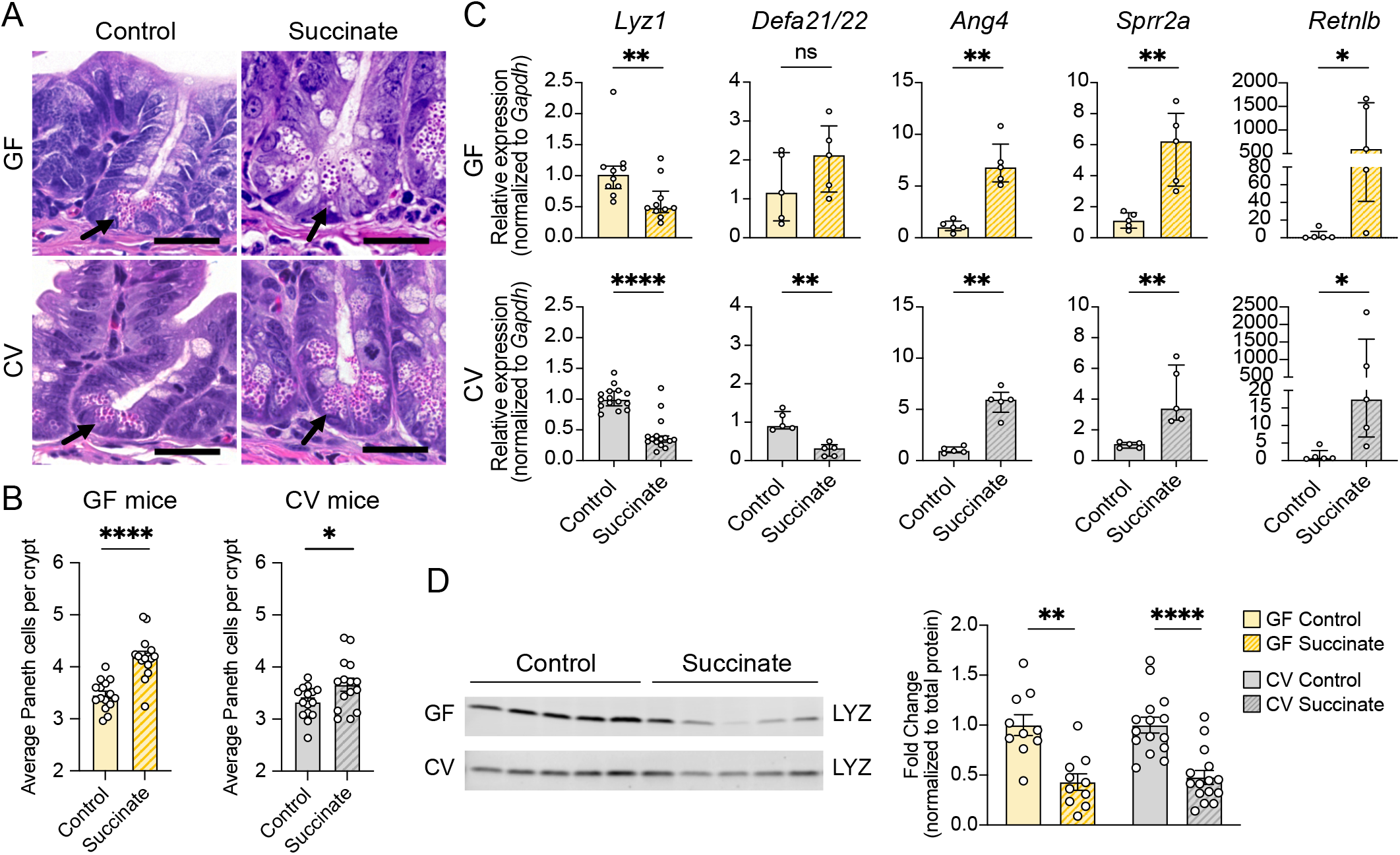
Oral administration of succinate results in similar changes to small intestinal AMP production as *T. mu* colonization. (A) Representative images of H&E-stained sections of the ileal crypts from control or succinate-treated GF (top row) or CV mice (bottom row). Scale bar: 25 μm. (B) Average number of Paneth cells per crypt in the ilea of control or succinate-treated GF or CV WT mice (n = 15 mice per group). Center values = arithmetic mean; error bars = SEM. Significance determined using Student’s *t*-tests. (C) Expression of representative AMP genes determined by RT-qPCR in the ilea of control or succinate-treated GF and CV mice (n = 5-15 mice per group). Relative expression normalized to *Gadph*. Center values = median; error bars = IQR. Significance determined using Mann-Whitney *U* tests. (D) Representative western blot images and quantitative analysis of intracellular lysozyme (LYZ) levels in the ilea of control or succinate-treated GF and CV mice. Each band or symbol represents an individual mouse. LYZ levels normalized to REVERT total protein stain (SI Appendix, Fig. S5E) (n = 10-15 mice per group). Center values = arithmetic mean; error bars = SEM. A linear mixed model was used to determine significance. ns = no significance, * *p* < 0.05, ** *p* < 0.01, *** *p* < 0.001, **** *p* < 0.0001.

Next, we assessed whether succinate alters antimicrobial expression. Succinate treatment increased *Dclk1* and *Muc2* expression (Fig. S5D) and altered expression of representative Paneth and goblet cell AMP genes in both GF and CV mice in a manner similar to *T. mu* colonization, increasing *Ang4, Sprr2a*, and *Retnlb* expression while decreasing *Lyz1* expression in both GF and CV mice (Fig. 5C). These transcriptional changes are consistent with decreased lysozyme protein levels (Fig. 5D and S5E-F), and increased RELMβ protein levels in response to succinate (Fig. S5F). While *Defa21/22* expression was decreased in succinate-treated CV mice, it was not significantly altered in succinate-treated GF mice in comparison to GF controls (Fig. 5C). Overall, these results show that succinate is sufficient to drive the same changes to AMP production that we observed in *T. mu-*colonized mice, independent of the resident microbiota.

To confirm that succinate sensing through the taste-chemosensory signal transduction pathway in tuft cells is critical for changes in AMP production, we treated *Trpm5*^-/-^ and *Sucnr1*^-/-^ mice with 100 mM succinate for one week in their drinking water. We found that lysozyme protein levels decreased in succinate-treated WT mice, but not in succinate-treated *Trpm5*^-/-^ and *Sucnr1*^-/-^ mice compared to their respective controls (Fig. S5G). Altogether, these data suggest that succinate is sufficient to remodel ileal epithelial cell composition and AMP production by stimulating tuft cell taste-chemosensory pathways.

Upon activation of taste-chemosensory signal transduction, SI tuft cells release IL-25, which induces ILC2s to produce type 2 cytokines such as IL-13 (20, 22–25). This cytokine signals to stem cells in the SI to bias their differentiation towards tuft and goblet cells. Additionally, helminth-induced type 2 immunity has been shown to alter expression of AMP-encoding genes in an IL-13-dependent manner (44, 45, 51). To determine if succinate-induced type 2 immunity and IL-13 are responsible for modulating ileal AMP production, we treated WT mice or mice lacking *Il13* (*Il13*^-/-^) with succinate for one week. Succinate induced tuft cell hyperplasia in WT mice, but not in *Il13*^-/-^ mice (Fig. S6A-B). In addition, Paneth cell morphology was unaffected by succinate treatment in *Il13*^-/-^ mice (Fig. S6C). Moreover, decreased *Lyz1* and *Defa21/22* expression, and increased *Ang4, Sprr2a*, and *Retnlb* expression found in succinate-treated WT mice were abrogated in succinate-treated *Il13*^-/-^ mice (Fig. 6A). Consistent with these results, we found no difference in intracellular lysozyme protein levels between control and succinate-treated *Il13*^-/-^ mice (Fig. S6D). These results indicate that changes to AMP production in response to succinate is dependent on type 2 immune induction, and specifically on the cytokine IL-13.

**Figure 6.**
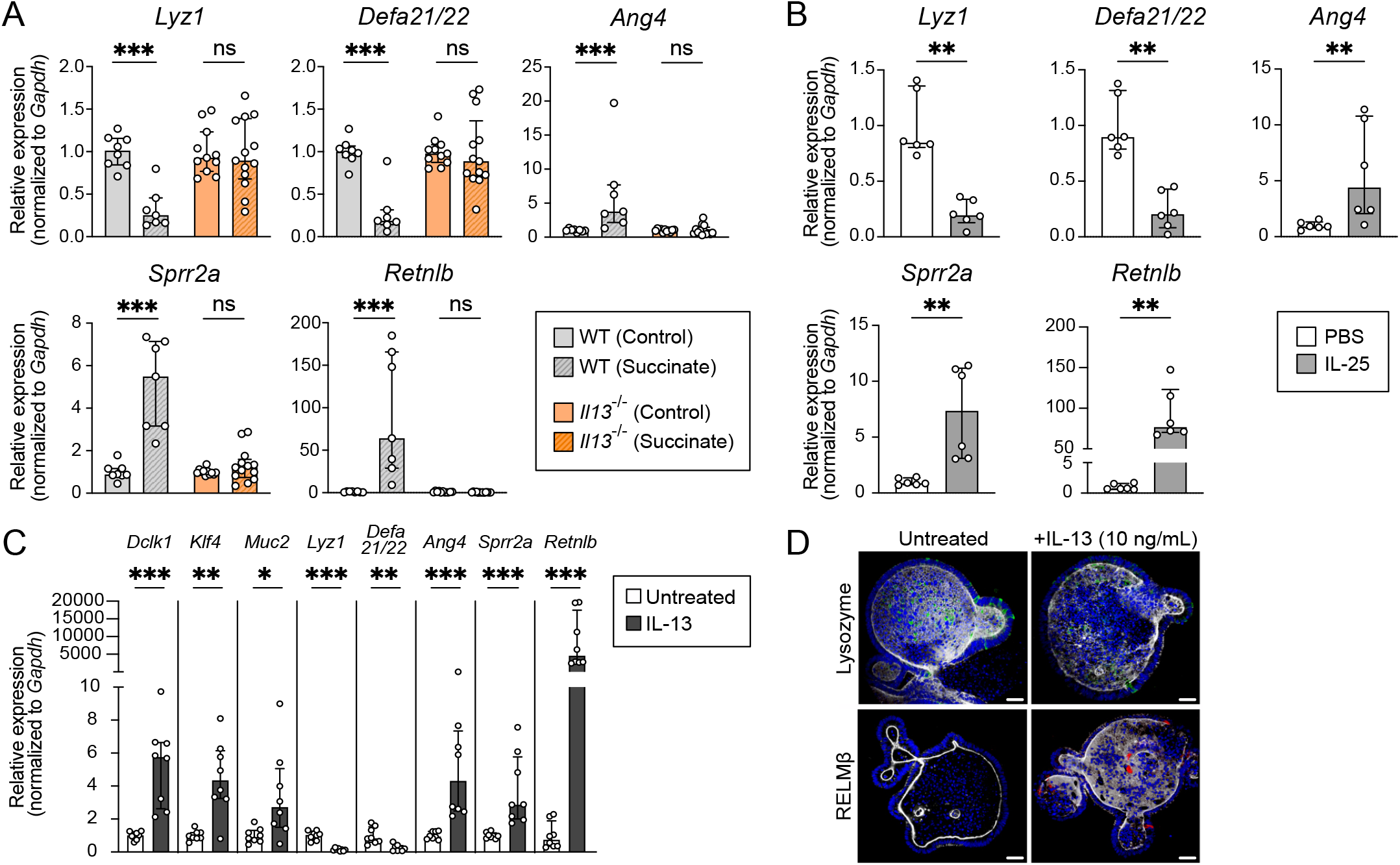
Succinate-induced changes to AMP production in the small intestine require type 2 immunity. (A) Expression of representative AMP genes determined by RT-qPCR in the ileal epithelial fraction of control or succinate-treated WT and *Il13*^-/-^ mice (n = 7-13 mice per group). (B) Expression of representative AMP genes determined by RT-qPCR in the ileal epithelial fraction of WT mice IP-injected with PBS or IL-25 (n = 6 mice per group). (C) Expression of *Dclk1* (tuft cell marker), *Klf4* and *Muc2* (goblet cell markers), and representative AMP genes determined by RT-qPCR in untreated or IL-13 treated ileal organoids (n = 8 samples per group). (D) Representative fluorescence microscopy images of untreated or IL-13 treated ileal organoids. Nuclei (blue), F-actin (white), Lysozyme (green), RELMβ (red). Scale bar: 30 μm. For RT-qPCR data, relative expression normalized to *Gapdh*. Center values = median; error bars = IQR. Significance determined using Mann-Whitney *U* tests. ns = no significance, * *p* < 0.05, ** *p* < 0.01, *** *p* < 0.001, **** *p* < 0.0001.

To determine if type 2 immune induction in the ileum is sufficient to alter host AMP production, we bypassed tuft cell stimulation and injected recombinant IL-25 intraperitoneally (IP) into WT mice (Fig. S6E). IL-25 injections induced tuft cell hyperplasia (Fig. S6F-G) and altered the expression of our representative panel of AMP genes in a similar manner as *T. mu* colonization or succinate administration (Fig. 6B). These AMP expression changes correlate with alterations in lysozyme organization within Paneth cells and an increase in RELMβ protein levels in IL-25 injected mice (Fig. S6H-I). This demonstrates that type 2 immune induction without initial tuft cell stimulation by succinate is sufficient to drive these changes to the SI epithelium.

To test if IL-13 directly stimulates remodeling of epithelial AMP expression, we used an *in vitro* primary intestinal organoid system (52, 53). SI organoids derived from mouse intestinal epithelial stem cells can differentiate into all epithelial subsets when cultured with specific growth factors and stimuli. IL-13 treatment of ileal organoids promoted tuft and goblet cell differentiation (20, 54), which we confirmed using RT-qPCR to measure the expression of the tuft cell marker *Dclk1* and goblet cell markers *Klf4* and *Muc2* by RT-qPCR (Fig. 6C). We also found that IL-13 treatment alone decreased epithelial expression of *Lyz1* and *Defa21/22*, while increasing expression of *Ang4, Sprr2a*, and *Retnlb* (Fig. 6C-D). Altogether, these results suggest that succinate stimulation of tuft cells leads to type 2 immune induction and IL-13 production, and that this cytokine remodels the distal SI epithelium and alters host AMP production.

### Type 2 immune induction alters the microbiome composition and depletes mucosa-associated microbes in the small intestine

AMPs play crucial roles in regulating the composition of the intestinal microbiota, controlling the extent of contact between microbes and the epithelium, and protecting the host from pathogen colonization (7, 33). Because succinate alters the expression of a wide range of AMP-encoding genes by stimulating type 2 immunity, we hypothesized that activation of type 2 immune responses would alter the composition of the ileal microbiota.

To induce type 2 immunity without confounding variables such as succinate feeding or *T. mu* colonization, which could independently alter the microbiota, we IP-injected IL-25 into mice every other day for 1 week (Fig. S6E and S7A), harvested the ileal luminal and mucosal microbial populations, and used qPCR to quantify the total number of bacterial 16S rRNA gene copies as a proxy for abundance. We saw no differences in 16S copy number between the luminal populations of IL-25-injected mice and mock-injected controls, indicating that luminal bacterial numbers were not altered by type 2 immunity (Fig. 7A). However, we observed significantly fewer 16S copies in the mucosal populations from IL-25-injected mice compared to controls (Fig. 7A), indicating that type 2 immune induction reduces the load of mucosa-associated bacteria.

**Figure 7.**
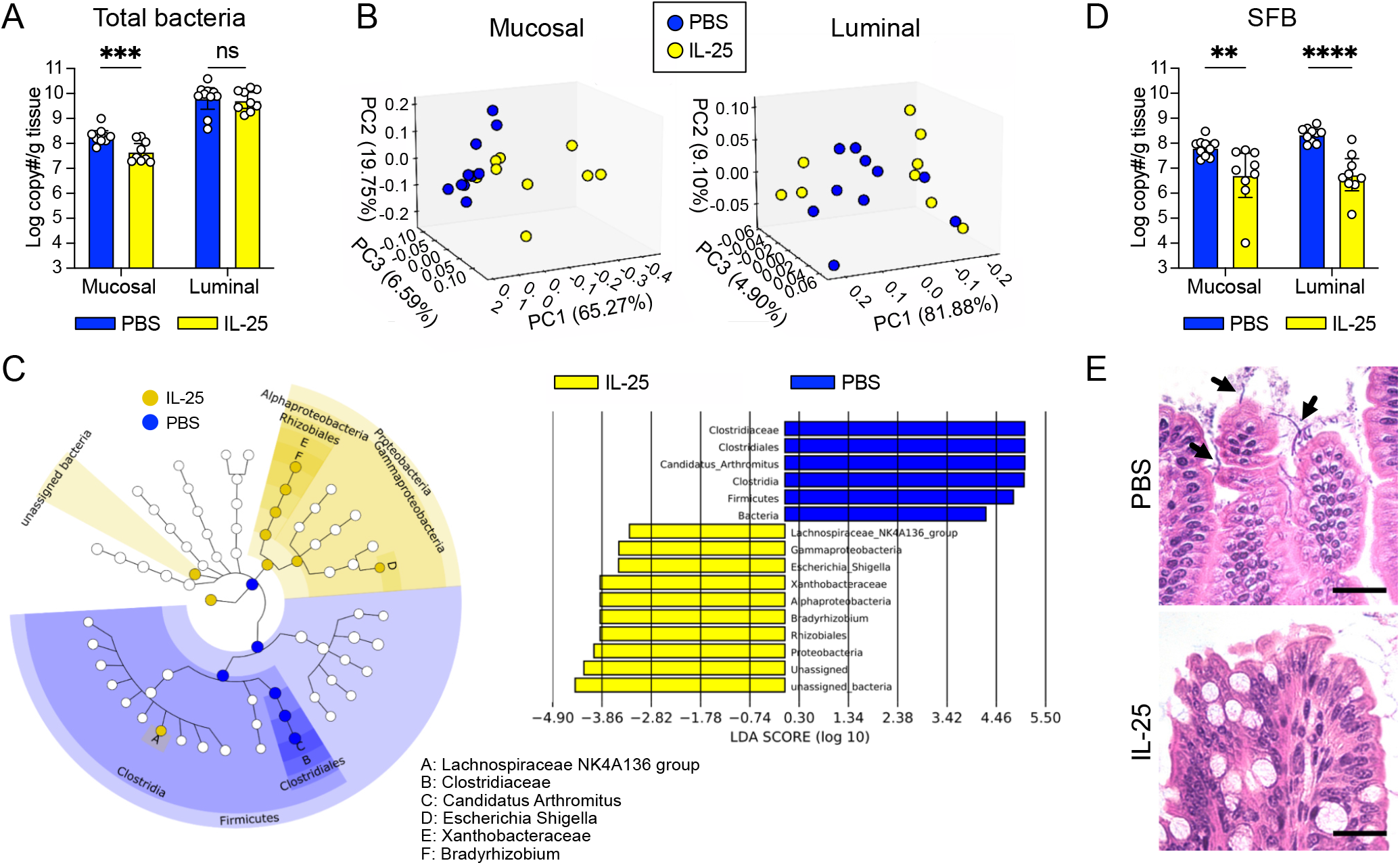
Type 2 immune induction depletes mucosa-associated bacteria in the small intestine. (A) Absolute quantification of total bacterial 16S rRNA gene copies in the ileal mucosal and luminal fractions of PBS or IL-25 injected mice. (B) Weighted UniFrac PCoA plots depicting differences in microbial communities in mucosal or luminal samples. The 1^st^, 2^nd^, and 3^rd^ principal components indicate the percentage of variation explained by the principal components for the ileal mucosal or luminal communities of PBS or IL-25 injected mice. Mucosal, *p* = 0.007; Luminal, *p* = 0.077. Significance determined via PERMANOVA test. (C) Linear discriminant analysis (LDA) effect size (LEfSe) analysis using a LDA threshold score of 2 to identify ileal mucosa-associated bacterial taxa in PBS or IL-25 injected mice. Cladogram (left) highlights taxonomic relatedness of bacteria, while LDA plot (right) is an ordered list of enriched bacteria. (D) Absolute quantification of SFB 16S rRNA gene copies in the ileal mucosal and luminal fractions of PBS or IL-25 injected mice. (E) Representative images of Carnoy’s-fixed, H&E-stained sections of ileal villi from PBS or IL-25 injected mice showing SFB (black arrows). Scale bar: 25 μm In Panels A and D, center values = geometric mean; error bars = 95% confidence interval. Significance determined using a generalized linear model. ns = no significance, * *p* < 0.05, ** *p* < 0.01, *** *p* < 0.001, **** *p* < 0.0001.

To obtain a broader view on how the composition of the ileal microbiota changes in the face of these AMP alterations, we conducted 16S rRNA gene sequencing analysis on these mucosal and luminal population samples. We performed β-diversity analysis to assess whether these communities are distinct from each other based on treatment conditions. Weighted UniFrac analysis, which accounts for both feature abundance and phylogenetic relatedness, showed that the mucosal populations of IL-25-injected mice are distinct from those of mock-injected mice (Fig. 7B). Altogether, these results suggest that type 2 immune induction by IL-25 injections is sufficient to alter the abundance and composition of the ileal mucosa-associated microbiota.

Next, we used Linear discriminant analysis Effect Size (LEfSe) analysis to determine which bacterial taxa are mostly likely to explain differences between the mock- and IL-25-injected mucosal populations. *Candidatus arthromitus*, commonly referred to as segmented filamentous bacteria (SFB), was one differentially abundant taxon in the mucosal populations of control mice (Fig. 7C). SFB adhere to the ileal epithelium (55, 56) and are the dominant members of mucosal communities in mice that harbor them (44). To confirm SFB depletion in the mucosa, we used qPCR to measure the number of SFB 16S copies in the mucosal and luminal samples as a proxy for SFB abundance. There were significantly less SFB 16S copies in both IL-25-injected mucosal and luminal samples in comparison to their respective controls (Fig. 7D); we further confirmed this by microscopy (Fig. 7E). Conversely, taxa that were enriched in the IL-25-injected mucosal populations include *Lachnospiraceae* and Proteobacteria (Fig. 7C). However, upon further examination, the Proteobacteria enriched in the IL-25 mucosal samples (i.e. *Bradyrhizobium, Escherichia*) are often found as contaminants in low biomass microbiota samples (57). Given that the total bacterial 16S copy number was reduced about 10-fold with IL-25 injections (Fig. 7A), these mucosal samples may have been susceptible to detecting these contaminants. We also performed LEfSe analysis on the luminal populations and found many unique taxa enriched in the mock-injected group (Fig. S7B). This suggests that despite no difference in the overall luminal bacterial numbers between both experimental groups (Fig. 7A), type 2 immunity may induce more subtle changes to the community composition. In addition, one of these enriched taxa is SFB, which is consistent with our observations of the mucosal populations in the mock-injected group. Altogether, these data show that SFB are differentially abundant between mock- and IL-25-injected mice in both luminal and mucosal microbiota fractions.

We also investigated whether type 2 immune-induced AMP alterations would decrease the abundance of other sensitive microbes, such as the Gram-positive human gut commensal *Enterococcus faecalis* (*Ef*). *Ef* is killed *in vitro* when incubated with purified recombinant ANG4 or SPRR2A (14, 44), AMPs that are induced with type 2 immunity (Fig. 6A-B). To assess the impact of type 2 immune-associated AMPs on *Ef* survival throughout the gastrointestinal (GI) tract, we first IP-injected mice with IL-25, and then orally inoculated these mice and mock-injected controls with *Ef* (Fig. S7C). The SI, cecum, and large intestine were collected from each mouse two hours post-gavage to enumerate *Ef* colony-forming units (CFUs). We found that *Ef* CFUs were significantly lower in the SIs of IL-25-injected mice compared to controls but not in other GI sites (Fig. S7C). This shows that type 2 immune induction lowers *Ef* abundance specifically in the SI, which could reflect an inhospitable AMP environment for this bacterium. However, given that type 2 immunity is typically associated with increases in gut motility (58), we assessed GI transit in mock- and IL-25-injected mice by repeating the above experiment, but administering fluorescent microspheres by oral gavage instead of *Ef*. At two hours post-gavage, luminal contents from the same GI regions were collected to quantify the number of microspheres present. We found equivalent microspheres in the SIs of mock- and IL-25-injected mice (Fig. S7D), suggesting that IL-25 injections at this dose and time frame do not alter gut motility within this region. Altogether, these results suggest that decreased *Ef* abundance in the SI of type 2 immune-induced mice is likely not due to increased peristalsis but may reflect increased *Ef* killing by AMPs.

## Discussion

Our results demonstrate that succinate produced by the commensal protist *T. mu* biases epithelial differentiation towards the secretory lineage and alters AMP expression in the distal SI. Succinate drives these changes by activating tuft cells, resulting in a type 2 immune response in the distal SI. Changes in AMP production in response to type 2 immunity alter the SI microbiota composition, with a specific reduction in the abundance of mucosa-associated bacteria such as SFB (Fig. S8) but not the abundance of protists (Fig. S4A). Previous studies have shown that oral treatment with polyethylene glycol (PEG) leads to mild diarrhea and a spike in luminal succinate due to the resulting bacterial dysbiosis (21, 59). Intriguingly, PEG-treated mice exhibited tuft cell hyperplasia specifically in the ileum (21), the SI region where tuft cells are most sensitive to succinate stimulation (22). These results indicate that short-term bacterial dysbiosis within the distal SI can increase luminal succinate concentrations and activate tuft cells. These observations, coupled with our study which revealed that commensal-derived succinate remodels AMP production and specifically reduced ileal mucosa-associated bacteria, suggest that tuft cell SUCNR1 may be attuned to sense bacterial-derived succinate rather than having evolved to detect protists in the distal SI.

Within the *T. mu*-colonized ileal epithelium, we saw an increase in ST spots containing tuft and goblet cell markers, which is indicative of their hyperplasia and consistent with prior studies (20, 22, 23). However, we also identified an increase in Paneth cell gene expression signatures in the *T. mu*-colonized ileum, which reflects a previously unknown response of this epithelial cell to commensal tritrichomonads. We then measured Paneth cell numbers within the crypts of uncolonized and *T. mu*-colonized mice and confirmed that Paneth cell hyperplasia was induced by protist colonization. Succinate, a metabolic byproduct of *T. mu*, was sufficient to increase Paneth cell numbers in the distal SI. In addition, *T. mu* colonization also altered Paneth cell secretory granule morphology and AMP localization. In uncolonized mice, Paneth cells typically contain large, electron-dense secretory granules surrounded by a thin, electron-lucent halo. Conversely, Paneth cells in *T. mu*-colonized mice contain smaller, electron-dense granules surrounded by larger, electron-lucent mucin globules. AMPs, such as lysozyme, are typically concentrated within the secretory granules at the apical side of Paneth cells. However, in *T. mu*-colonized mice, lysozyme was dispersed throughout the cell. This altered granule morphology and lysozyme distribution is reminiscent of Paneth cells observed in mice with mutations in susceptibility genes linked to Crohn’s disease in humans (60–62). *Atg16L1* encodes an autophagy protein, and the loss of this gene in mice led to abnormal lysozyme distribution, indicating defects in packaging AMPs into secretory granules (60). Furthermore, mutations in *Atg16L1* disrupt the release of AMPs through the secretory autophagy pathway during bacterial infection (63). These observations support the idea that *T. mu* colonization may alter Paneth cell autophagy, potentially affecting their secretory function. However, type 2 immune induction significantly reduced mucosa-associated bacteria, suggesting that these changes to Paneth cell secretory granule morphology and AMP localization do not interfere with AMP secretion or microbial killing. Determining how succinate and type 2 immunity influences Paneth cell functions and AMP secretion will be important for future investigation.

We also observed that *T. mu* profoundly shifted ileal AMP expression and production in the distal SI. Intriguingly, these transcriptional changes did not solely consist of upregulation of all AMP-encoding genes, as one might expect with Paneth cell hyperplasia. AMPs such as *Reg3b, Reg3g, Lyz1*, and *Defa21/22* dramatically decreased, indicating that the host selectively remodeled the antimicrobial landscape rather than uniformly increasing AMP production. Succinate was sufficient to elicit these same changes in AMP expression. While commensal microbes (12–15, 42, 46) and helminth infections (44, 45, 51) can alter AMP expression, these previous studies did not implicate microbial metabolites in this process. However, helminths and *T. mu* both induce intestinal type 2 immunity, resulting in similar downregulation of the AMPs *Lyz1, Defa21/22*, and *Reg3g*, and upregulation of *Ang4, Sprr2a*, and *Retnlb* (44, 51). Despite both organisms stimulating type 2 immune responses and IL-13 production, they affect different locations in the SI. The helminth *Heligmosomoides polygyrus* infects the proximal SI (duodenum and proximal jejunum), where expression of AMPs such as *Sprr2a* were most profoundly altered (44). In contrast, *T. mu* colonizes the distal SI (distal jejunum and ileum), which contains 3- to 7-fold more Paneth cells than the proximal SI and is adjacent to densely-populated bacterial communities in the cecum and colon (64, 65). In addition, the helminth ligand responsible for inducing type 2 immunity is still unknown; proximal SI tuft cells are relatively insensitive to succinate and SUCNR1 is not required for initiating anti-parasitic responses (21, 22). Thus, there may be an important biogeographical distinction between how AMPs are affected by pathogenic worms and commensal protists.

AMPs are critical mediators of host-bacterial interactions, leading us to investigate how type 2 immunity impacted the overall numbers and composition of the ileal luminal and mucosal microbiota. We found that type 2 immune induction significantly decreased the total number of mucosa-associated bacteria, while the luminal population largely remained unaffected. This was surprising because *T. mu* colonization downregulates *Reg3g*, an AMP that is known to be important for segregating commensals from the gut epithelium (5). Therefore, we hypothesize that other AMPs upregulated by type 2 immunity can compensate for reduced *Reg3g* and enhance the killing of mucosal microbes. Indeed, compared to controls, the absolute abundance of SFB is reduced in the mucosal populations of IL-25 injected mice. SFB are often the dominant taxa at the ileal mucosa (44), but previous studies showed that SFB abundance can be reduced by mucosal immunity and AMPs. For instance, SFB numbers decrease during helminth infection (51) or in mice transgenic for a human Paneth cell defensin (9), while SFB numbers increase in *Sprr2a*-deficient mice (44). These observations are consistent with our results, as IL-25 injections upregulate the AMP *Sprr2a* and reduce SFB abundance simultaneously. Collectively, this data suggests that type 2 immune induction can restructure the bacterial microbiota composition. AMPs are likely involved in these population changes, as these peptides are embedded within the mucus layer close to the mucosal surface (4).

Succinate is produced as a metabolic byproduct of redox homeostasis in protists belonging to the class *Tritrichomonadea* (66), but colonization abundance by tritrichomonads like *T. mu* is not affected by succinate-driven type 2 immunity (20, 22). Furthermore, *T. mu* does not damage tissue during chronic colonization like parasitic helminths, implying that fortification of the intestinal barrier by increasing mucus production and altering AMP expression is for a different purpose. Instead, our findings suggest that tuft cell sensing of succinate and the subsequent AMP expression changes may have evolved to respond to bacterial disturbances. Succinate is a common metabolic byproduct produced by commensal bacteria during carbohydrate fermentation. However, it is present at relatively low concentrations in the gut lumen due to rapid consumption by other microbes (26, 27). Thus, succinate accumulation in the gut lumen is generally considered abnormal and associated with dysbiotic events such as antibiotic treatment and gut motility disturbances (21, 59). In addition, the loss of goblet and Paneth cells leads to a bloom in succinate-producing bacteria, further reinforcing the connection between secretory epithelial cells and this metabolite (29). Thus, it is possible that tuft cells have evolved to respond to increased luminal succinate secondary to bacterial dysbiosis, triggering type 2 immunity and altering AMP production to restructure the bacterial microbiota and restore homeostasis.

## Author Contributions

C.F., L.M.F., M.R.H., and N.H.S. designed research. C.F., L.M.F., and M.B.G. performed and analyzed the research. G.M.B. prepared tissues for spatial transcriptomics and conducted transcriptomic data analysis. S.N.A. conducted 16S rRNA sequencing data analysis. E.R.G. provided help with protist culture/colonization and flow cytometry. M.H., J.A.L., K.F.N., M.D.W.T., and L.B.D. provided help with sample harvests and data collection. J.Z. provided help with germ-free mouse colony maintenance and experiments. A.P. conducted statistical analyses of experimental data. C.F., L.M.F., G.M.B., S.N.A., M.R.H., and N.H.S. wrote the manuscript with input from all authors.

## Acknowledgments

We thank the Howitt Lab, Salzman Lab, and Dr. K. Ng for comments on the manuscript and helpful discussions. We thank Dr. R. Margolskee, Amgen, and Dr. C-H. Lee for generously sharing the *Trpm5*^-/-^ mice, *Sucnr1*^-/-^ mice, and *Il13*^-/-^ mice, respectively. We thank C. Mechler for her help with mouse colony maintenance and genotyping. We thank A. Torres and H. Tang for their help with cryosectioning tissue. We thank P. Chu at Stanford’s Pathology Department Histology Service Center and the Core Histology Facility at the Children’s Research Institute (CRI), Children’s Wisconsin for processing tissue samples, and Dr. S. Kumar at the CRI Core Imaging Facility for scanning microscopy slides. We thank J. Perrino at the Stanford Cell Sciences Imaging Facility (CSIF) for transmission electron microscopy (TEM). We thank Dr. D. Wagh and Dr. J. Coller from the Stanford Functional Genomics Facility (SFGF) for their help with spatial transcriptomics (ST). We thank the UW-Madison Biotechnology Center for 16S rRNA sequencing. Cartoon schematics were created with BioRender.com, except for Fig. S8.

This research was supported by the Stanford School of Medicine Dean’s Postdoctoral Fellowship, Stanford Maternal & Child Health Research Institute (MCHRI) Postdoctoral Fellowship, and A.P. Giannini Foundation Postdoctoral Fellowship (to C.F.); National Institutes of Health (NIH) R01DK128292 and R21AI171222 (to M.R.H.); the Medical College of Wisconsin’s Digestive Disease Center, the Advancing a Healthier Wisconsin Endowment, and NIH R35GM122503 (to N.H.S.); NIH T32 AI00729037 (to G.M.B. and K.F.N.); the Stanford MCHRI Pediatric IBD and Celiac Disease Research Postdoctoral Fellowship (to E.R.G.); the NSF Graduate Research Fellowship and Stanford Graduate Fellowship (to M.D.W.T.); and the Stanford Bio-X Undergraduate Summer Research Program Award (to L.B.D.). ST sequencing data was generated with instrumentation at the SFGF, which were purchased with NIH awards S10OD025212 and 1S10OD021763. TEM done by Stanford’s CSIF was supported, in part, by NIH S10 Award Number 1S10OD028536-01 from the Office of Research Infrastructure Programs (ORIP). The data/contents are solely the responsibility of the authors and do not necessarily represent the official views of the National Center for Research Resources (NCRR) or the NIH.

## Materials and Methods

### Animal work ethics statement

All animal experiments were performed in accordance with NIH guidelines, with approval by the Institutional Animal Care and Use Committees (IACUC) of Stanford University and the Medical College of Wisconsin. Animals were housed in research animal facilities at Stanford University or the Medical College of Wisconsin that are accredited by the Association of Assessment and Accreditation of Laboratory Animal Care (AAALAC) International.

### Mouse strains and husbandry

WT C57BL/6J mice (Stock #000664) were purchased from Jackson Laboratory. C57BL/6 mice *Gfi1b*^EGFP/+^ mice were purchased from Jackson Laboratory (Stock #016161) and were bred in-house at Stanford University. C57BL/6 *Trpm5*^-/-^ mice were generously provided by Dr. Robert Margolskee at the Monell Chemical Senses Center (49).

C57BL/6 *Sucnr1*^-/-^ mice were generously provided by Amgen under a materials transfer agreement. C57BL/6 *Il13*^-/-^ mice were generously provided by Dr. Chi-Hao Lee at the Harvard School of Public Health (67). *Sucnr1*^-/-^ mice were bred in-house at Stanford University, and experiments were performed using co-housed WT littermates. *Trpm5*^-/-^ and *Il13*^-/-^ mice were bred in-house at Stanford University, and co-housed with WT controls for 2 weeks prior to use in experiments. For germ-free experiments, WT C57BL/6J mice were bred and maintained in semi-rigid isolators (Park Bioservices) under standard germ-free conditions in the Gnotobiotic Core Facility at the Medical College of Wisconsin. For 16S rRNA sequencing experiments, WT C57BL/6 male mice (5 weeks old; Model # B6-M MPF) were purchased from Taconic Biosciences (San Diego, CA), whose mouse colonies naturally contain segmented filamentous bacteria (SFB) as part of their microbiome (56).

### Isolation of *Tritrichomonas musculis* for mouse colonization

A colony of WT C57BL/6J mice colonized with *T. musculis* is maintained at Stanford University as a protist source. To obtain protists for colonization experiments, *T. musculis* was isolated from source animals and cultured overnight at 37°C in an anaerobic cabinet (Coy Lab Products), as previously described (20, 68). The following day, 1×10^5^ *T. mu* were orally administered to each mouse. *T. musculis*-colonized mice were sacrificed 3 weeks post-colonization.

### 10x Visium Spatial Transcriptomics (ST)

The distal 5 cm of the small intestine from uncolonized and *T. musculis*-colonized mice were harvested. The luminal contents were flushed out with ice-cold sterile PBS using a 19-gauge feeding needle, and the tissue was opened longitudinally and swiss-rolled. The swiss-rolled tissue was then covered in OCT (VWR) and immediately frozen in an isopentane (Sigma-Aldrich) and liquid nitrogen bath. The frozen tissues were stored at -80°C until further processing.

10 μm-thick sections were cut and placed onto the Visium Gateway slide (10x Genomics, Pleasanton, CA) to perform gene expression analysis. The manufacturer’s protocol was followed without any significant alterations. Briefly, sections were fixed with methanol for 30 minutes at -20°C, followed by hematoxylin and eosin (H&E) staining and imaging using a BZ X-800 microscope (Keyence, Itasca, IL). Sections were subjected to tissue permeabilization for 10 minutes—permeabilization time was established using a Tissue Optimization slide (10x Genomics) and the manufacturer’s protocol. This was followed by reverse transcription for cDNA synthesis, second strand synthesis, cDNA amplification (18 cycles), and NGS library preparation. Libraries were quantified using Bioanalyzer (Agilent, Santa Clara, CA) and quantitative PCR (qPCR) analysis (Bio-Rad, Hercules, CA). Libraries were pooled and sequenced on the NovaSeq 6000 instrument (Illumina Inc., San Diego, CA), and data was processed through the Space Ranger pipeline (10x Genomics).

### Filtering and normalization of small intestinal spatial transcriptomics (ST) data

All ST data analysis was computed in R. ST spots with less than 300 genes detected or more than 50% mitochondrial gene expression were first filtered out as low-quality cells and were excluded from the analysis. The FindVariableFeatures function was used to select 2,000 variable genes with default parameters. The ScaleData function was used to scale and center the counts in the dataset. STutility was used to process gene expression matrices for the uncolonized and *T. musculis*-colonized tissue sections using the InputFromTable function. Effectively, this merged and converted the samples into a single Seurat object. The merged dataset was then enriched for protein coding genes by removing genes annotated with non-coding RNA. The filtered dataset was then normalized by variance stabilizing transformation using Seurat’s SCTransform.

### Visualization of spatial expression

STUtility (30) offers a variety of native tools to visualize the expression of genes on the tissue. We took advantage of the FeatureOverlay function to visualize our genes of interest. Moreover, the VlnPlot function in Seurat was used to visualize relative changes in gene expression between the samples.

### Deconvolution of ST data

Using non-negative matrix factorization (NNMF), the spatially-resolved transcriptomic data set was split into the respective uncolonized and *T. musculis*-colonized samples and deconvolved into 20 factors. A second mode of analysis included integrating the two sample datasets and deconvolving 20 factors resulting in gene modules and cell topics that described the data. Using the NNMF modes as the reduction, Harmony was used to integrate the uncolonized and *T. musculis*-colonized sample using the SCT assay values. 8 clusters were created and further defined by the top 5 genes with the largest log-2 fold change. Differential expression analysis (DEA) was conducted using FindAllMarkers from Seurat to detect marker genes for each cluster.

Moreover, we leveraged an existing small intestinal epithelial single-cell RNA sequencing (scRNA-seq) dataset (32) to map onto our ST dataset using the FindTransferAnchors within Seurat. This function constructs a weight matrix that defines the association between each query cell in the reference dataset and each anchor in the spatial dataset. These weights sum to 1 and were used as the percentage of the cell type in the spots. This results in a predicted ID score for the probability of the annotated cells present within the ST dataset. We used these predicted ID scores to discern the presence of certain epithelial cells on our tissue in an unbiased way.

### *Tritrichomonas musculis* enumeration

*T. musculis* numbers in the distal small intestine were enumerated as previously described (20, 68). Briefly, the distal 10 cm of the small intestine (ileum) was removed and flushed with ice-cold sterile PBS using a 19-gauge feeding needle.

The intestinal contents were pelleted by centrifugation at 2000x*g* for 5 minutes and stored at - 20ºC. Genomic DNA was isolated from the intestinal contents with the DNeasy PowerSoil Kit (Qiagen) according to the manufacturer’s instructions. To detect and enumerate *T. musculis*, qPCR was performed using the PowerUp SYBR Green Master Mix (Applied Biosystems) with the following primers recognizing *the T. musculis* 28S rRNA gene: *T. musculis* 28S rRNA for and *T. musculis* 28S rRNA rev (see Table S1 for primer sequences). To convert qPCR Ct values into protist numbers, we used a previously published standard curve that was generated using known amounts of *T. musculis* (20).

### Epithelial cell isolation and flow cytometry

For most experiments in this study, the distal 10 cm of the small intestine (ileum) was removed and flushed with ice-cold sterile PBS using a 19-gauge feeding needle. For experiments that involved the proximal small intestinal regions, the proximal 7 cm of the small intestine was removed for the duodenum, and the next 10 cm following that was removed for the jejunum and similarly flushed. The tissues were then opened longitudinally, Peyer’s patches were removed, and the remaining tissue was gently agitated at 4ºC in PBS, 2% heat-inactivated FBS (iFBS), 5 mM HEPES, and 1 mM DTT for 10 minutes. Then, the tissue was transferred into pre-warmed PBS with 2% iFBS, 5 mM HEPES, and 5 mM EDTA and shaken at 37ºC for 15 minutes, followed by vigorous shaking to strip off the epithelial cells. This was repeated once more, and epithelial cells from both fractions were combined and washed with PBS. Afterwards, the epithelium was digested in DMEM containing 10% iFBS, 0.5 U/mL Dispase II (StemCell Technologies), and 50 μg/mL DNase (Alfa Aesar) for 10 minutes at 37ºC. The resulting solution was passed through 70 μm filters and washed in PBS containing 2% iFBS and 1 mM EDTA. The resulting single-cell suspension was blocked with anti-CD16/CD32 (Clone 93, BioLegend), and then stained with the following antibodies: PacBlue-conjugated anti-CD45 (Clone 30-F11, BioLegend), Phycoerythrin/Cyanine7 (PE/Cy7)-conjugated anti-Ep-CAM (Clone G8.8, BioLegend), and AlexaFluor 647-conjugated anti-Siglec-F (Clone E50-2440, BD Biosciences). Cell viability was assessed by propidium iodide (BioLegend) staining. Cells were analyzed on a BD FACS Canto II. Epithelial cells were gated on viable CD45^-^ EpCAM^+^ cells. Tuft cells were further selected as SiglecF^+^ (gating strategy shown in Fig. S9). For experiments that involved *Gfi1b*^EGFP/+^ mice, in which tuft cells are specifically marked with EGFP, the single-cell suspension was stained with the following antibodies: PacBlue-conjugated anti-CD45 (Clone 30-F11, BioLegend) and APC-conjugated anti-EpCAM (Clone G8.8, BioLegend). Epithelial cells were gated on viable CD45^-^EpCAM^+^ cells. Tuft cells were further selected as EGFP^+^. Information on flow reagents is listed in Table S2.

### Transmission electron microscopy

The distal small intestines from uncolonized or *T. musculis*-colonized mice were harvested. A 1 cm long piece of the most distal part of the intestinal tube was cut and immersed in Karnovsky’s fixative—2% glutaraldehyde and 4% paraformaldehyde in a 0.1M sodium cacodylate buffer, pH 7.4, at room temperature (RT). After the intestinal tubes had been immersed for about 15 minutes, a scalpel was used to cut 1-3 mm rings from the intestinal tubes. These intestinal rings were fixed for an additional hour at RT, and then submitted to the Cell Sciences Imaging Facility at Stanford University for further sample processing. The fix was replaced with cold-aqueous 1% osmium tetroxide and then allowed to warm to RT for 2 hours rotating in a hood, washed 3X with ultrafiltered water, then en bloc stained in 1% uranyl acetate at RT for 2 hours while rotating. Samples were then dehydrated in a series of ethanol washes for 30 minutes each at RT beginning at 50%, then 70% ethanol, and then moved to 4 ºC overnight. Afterwards, the samples were placed in cold 95% ethanol and allowed to warm to RT, change to 100% ethanol 2X, and then propylene oxide (PO) for 15 minutes. Samples were infiltrated with Embed-812 mixed 1:2, 1:1, and 2:1 with PO for 2 hours each, with samples left in 2:1 resin to PO overnight, rotating at RT in a hood. The samples are then placed into Embed-812 for 2 to 4 hours, placed into molds with labels and fresh resin, oriented, and placed into a 65ºC oven overnight. Sections were taken around 80 nm using an UC7 (Leica, Wetzlar, Germany), picked up on formvar/Carbon-coated 100 mesh Cu grids, stained for 40 seconds in 3.5% uranyl acetate in 50% acetone, followed by staining in Sato’s Lead Citrate for 2 minutes. Samples were observed in the JEOL JEM-1400 120kV electron microscope. Images were taken using a Gatan Orius 832 rk X 2.6k digital camera with 9 μm pixels.

### Histology and fluorescence microscopy

The distal small intestine was removed and fixed in 4% PFA diluted in PBS at 4°C overnight, zinc formalin at room temperature overnight, or Carnoy’s solution at room temperature overnight. The tissue was then embedded in paraffin and cut into 4 μm-thick sections. H&E or Alcian blue staining was performed using standard procedures by the Stanford Pathology Department Histology Service Center or the Core Histology Facility at the Children’s Research Institute (CRI), Children’s Wisconsin. For immunofluorescence, heat-mediated antigen retrieval was performed in sodium citrate buffer (10 mM sodium citrate, 0.05% Tween 20, pH 6.0) for 20 minutes. Then, the slides were washed in PBS and blocked in PBS containing 3% BSA, 2% goat serum, and 0.1% saponin for 1 hour at room temperature. Primary antibodies were incubated at 4ºC overnight, and secondary antibodies were incubated for 1 hour at room temperature. Primary antibodies used include: Rabbit anti-DCLK1 (1:200 dilution, Abcam), Mouse anti-E-Cadherin (1:400 dilution, BD Biosciences), Rabbit anti-Lysozyme (1:500 dilution, Agilent), Rabbit anti-RELMβ (1:200 dilution, PeproTech), Rabbit anti-MUC2 (1:200 dilution, Abcam), and DNA was labeled with DAPI (0.5 μg/mL). Samples were washed and then mounted in VECTASHIELD Antifade Mounting Medium (Vector Laboratories). Images were captured with a Zeiss LSM 700 confocal microscope (Carl Zeiss, Oberkochen, Germany) or a Nikon Eclipse E400 microscope (Nikon). Information on antibodies used is listed in Table S2.

### Paneth cell enumeration

Paneth cells were identified using H&E-stained slides. For Paneth cell counts, only crypts that were aligned along the longitudinal axis such that the crypt lumen could be seen were evaluated. Granule-containing Paneth cells per crypt were manually counted on a Nikon Eclipse E400 microscope (Nikon) or from scanned slide images. Slides were scanned and digitized by the Imaging Core at the Children’s Research Institute, Children’s Wisconsin using a Hamamatsu slide scanner or by the Stanford Pathology Department Histology Service Center using an Aperio AT2 slide scanner (Leica). Image analysis of scanned slides was conducted manually using OlyVIA (Olympus). Paneth cell enumeration was represented as the average number of Paneth cells per crypt.

### Tuft cell enumeration via immunofluorescence microscopy

Tuft cells were identified by immunofluorescence microscopy using DCLK1 staining as described above. Images were acquired using a VS120 slide scanner (Olympus) at the Imaging Core at the Children’s Research Institute, Children’s Wisconsin. Image analysis was conducted manually using QuPath. First, the polyline tool was used to measure 20 crypt-villus axis lengths per ileum, using the DAPI and E-Cadherin channels as reference. Next, the DCLK1 channel was used as reference to enumerate tuft cells among the previously established regions of interest. Tuft cell enumeration was represented as the number of tuft cells per mm crypt-villus axis.

### RNA isolation and reverse transcription-quantitative PCR (RT-qPCR) for *in vivo* and *in vitro* samples

For RNA isolation from total epithelium from the distal small intestine (samples from Fig. 4 and 6), the epithelial fraction was collected and lysed in Qiazol (Qiagen) for RNA extraction according to the manufacturer’s instructions. For small intestinal organoids, whole organoids were lysed in Qiazol, and RNA was extracted accordingly. RNA concentrations were determined using a NanoDrop. RNA was DNase-treated using the TURBO DNA-free Kit (ThermoFisher Scientific), and then cDNA was synthesized using the iScript cDNA Synthesis Kit (Bio-Rad). RT-qPCR was performed using the PowerUp SYBR Green Master Mix (Applied Biosystems).

For samples from Fig. 5, freshly collected or RNAlater (ThermoFisher Scientific)-preserved sections of the distal small intestine were manually homogenized using a pestle tissue grinder assembly and RNA was extracted using a RNeasy minikit (Qiagen) following the manufacturer’s instructions. RNA was DNase-treated using the TURBO DNA-free kit (ThermoFisher Scientific). RNA was further purified using sodium acetate precipitation. RNA concentrations were determined using a NanoDrop, and cDNA synthesis was performed using the iScript cDNA Synthesis Kit (Bio-Rad). RT-qPCR was performed using the iTaq Universal SYBR Green Supermix (Bio-Rad). All RT-qPCR primer sequences are listed in Table S1.

### Immunoblotting

The distal small intestine was removed and flushed with ice-cold sterile PBS using a 19-gauge feeding needle. The ilea were opened longitudinally and then snap-frozen in liquid nitrogen. When ready for processing, the ilea were thawed and homogenized in 2 mL of RIPA buffer and 20 μL HALT protease inhibitor cocktail (100x) using a handheld electronic tissue homogenizer. The homogenates were then incubated on a shaker at 4ºC for 20 minutes to extract intracellular proteins. The homogenates were pelleted by centrifugation at 16000xg for 15 minutes at 4ºC. The supernatants, which contain extracted proteins, were transferred to new tubes and pelleted once again at the same settings to remove any residual debris. Protein concentrations for each sample were determined using the Pierce BCA Assay Kit. Equivalent amounts of extracted intracellular proteins (10-20 μg) were separated by SDS-PAGE and transferred onto PVDF membrane. The PVDF membrane was blocked in LI-COR Intercept blocking buffer (LI-COR Biosciences) for one hour, and lysozyme was detected using a Rabbit anti-Lysozyme antibody (1:10000 dilution, Abcam), followed by an IRDye® 800CW Goat anti-rabbit IgG (H+L) or IRDye® 800CW Donkey anti-rabbit IgG (H+L) secondary antibody (LI-COR Biosciences). The blot was then scanned and imaged on a LI-COR Odyssey Imager. REVERT total protein stain was used for normalization of lysozyme levels. Information on antibodies used is listed in Table S2.

### Succinate feeding

For succinate treatment, mice were provided with 100 mM succinic acid disodium salt (Sigma-Aldrich; Cat# 224731-500G) in their drinking water for 1 week. 200 mM sodium chloride (Sigma-Aldrich; Cat# S9888-500G) was used as a control to match sodium molarity, except for experiments with germ-free (GF) and conventional (CV) mice in Fig. 5 and Fig. S5A-E, where water was used as a control.

### *In vivo* IL-25 injections

WT mice were intraperitoneally (IP)-injected every other day with 500 ng of recombinant IL-25 (R&D Systems; Cat# 7909-IL-010/CF) or equivalent volume of sterile PBS (vehicle control) over the course of one week. The distal small intestine was harvested to examine the abundance of tuft cells, expression of tuft cell and Paneth cell markers, and Paneth cell morphology as described above.

### Small intestinal organoid culture

Distal small intestinal organoids were prepared as previously described (20, 52, 69). When indicated, IL-13 (10 ng/mL) (BioLegend; Cat# 575902) was added to L-WRN growth media for 2 days.

### Small intestinal organoid imaging

Small intestinal organoids embedded in Matrigel were seeded onto circular glass coverslips, allowed to grow, and then treated with the desired conditions. Afterwards, the samples were fixed in 2% PFA at 4°C overnight. The samples were washed in PBS 3x the next morning and stored in PBS at 4°C until further processing. For immunofluorescence, organoids were incubated with primary antibodies diluted into PBS containing 3% BSA and 0.2% Triton X-100 overnight at room temperature. Next morning, the samples were washed in PBS 3x, and secondary antibodies were incubated for 3 hours at room temperature. Samples were washed in PBS 3x and then mounted in VECTASHIELD Antifade Mounting Medium (Vector Laboratories). Primary antibodies used include: Rabbit anti-Lysozyme (1:100 dilution, Agilent) and Rabbit anti-RELMβ (1:100 dilution, PeproTech). DNA was labeled with DAPI (0.5 μg/mL) and F-actin was labeled with AlexaFluor™ 660 Phalloidin (ThermoFisher Scientific). Images were captured with a Zeiss LSM 700 confocal microscope (Carl Zeiss, Oberkochen, Germany). Information on antibodies used is listed in Table S2.

### Total bacteria and segmented filamentous bacteria (SFB) 16S copy number enumeration

To quantify the number of total bacterial or SFB 16S rRNA gene copies in the ileal mucosal and luminal fractions from the DNA samples used for 16S rRNA sequencing, qPCR was performed using the iTaq Universal SYBR Green supermix (Bio-Rad) with primers recognizing total bacteria (UniF340 and UniR514) or SFB (SFB736F and SFB8444R) (see Table S1 for primer sequences). Absolute quantification of 16S rRNA copies was determined using standard curves that were constructed using previously published plasmids that contain 16S rRNA genes specific to each group analyzed (total bacteria or SFB) (70). To track total bacterial and SFB 16S copy number prior to and after IL-25 injections, we collected fecal pellets from the animals at the said timepoints, isolated genomic DNA from the feces, and used the same procedure as described above to detect and enumerate total bacterial and SFB 16S copies.

### Sample processing and DNA extractions for 16S rRNA sequencing

WT Taconic mice were IP-injected with PBS or IL-25 as described above. At the experimental endpoint, the distal 10 cm of the small intestine was harvested from both mouse groups. The terminal 1 cm was removed for histological analysis, and the remaining 9 cm were separated into mucosal and luminal fractions. For isolation of luminal contents, the ileum was flushed with ice-cold sterile PBS using a 19-gauge feeding needle, and then pelleted by centrifugation at 2000x*g* for 5 minutes. The supernatants were removed, and the luminal contents were saved for further processing. The remaining ileal tissue was used for the mucosal fraction. Genomic DNA was extracted from the mucosal and luminal fractions using the DNeasy PowerSoil Kit (Qiagen). DNA was sent to UW-Madison Biotechnology Center for 16S sequencing. The V3/V4 region (341-806) of the 16S rRNA gene was amplified by PCR (341F_primer: CCTACGGGNGGCWGCAG, 806R_primer: GACTACHVGGGTATCTAATCC) and sequenced on the MiSeq platform (Illumina) using 2 × 300 base pair paired-end protocol, which yielded paired-end reads.

### 16S rRNA sequencing data analysis

QIIME2 (v. 2022.2) was used to analyze the paired-end 16S rDNA sequencing reads (71). Sequences were imported and summarized to check quality. Cutadapt was used to trim primers from the reads (72). Representative sequences were chosen using DADA2, which also removes chimeric sequences (73). The representative sequences were then aligned (74), masked for hypervariable regions (75), and phylogenetic trees were produced (76). A *classifier* was generated to assign taxonomy to the reads using the 99% similarity files of the SILVA v. 138 and the 341-806 region (V3/V4) of the 16S gene (77). Taxonomy was assigned to the feature table to make taxonomy bar plots and to generate relative abundance tables. Diversity metrics were run using the *core-metrics-phylogenetic* command of QIIME2. Alpha and beta diversity were analyzed using their respective commands, *alpha/beta-group-significance* (78–80). Alpha diversity metrics used a Kruskal-Wallis test to test for significance, while beta diversity metrics used a PERMANOVA test to test for significance; both types of metrics used Benjamini-Hochberg multiple comparison tests. Principal Coordinate Analysis (PCoA) plots were examined using EMPeror (81, 82) and finalized figures were made using Python (83). LEfSe, Linear discriminant analysis (LDA) effect size, was run to determine enriched organisms for each treatment group (84). Final LEfSe figures were generated using Inkscape v. 1.0.1 (https://inkscape.org/). Cladograms were generated using GraPhlAn (85).

### *Enterococcus faecalis* culture

*E. faecalis* strain OG1RF (86) was cultured aerobically at 37°C in Mueller-Hinton (MH) broth (BD Life Sciences) supplemented with 50 μg/mL rifampin (Rif) (Chem-Impex Int’l Inc.), or on MH agar plates supplemented with 200 μg/mL Rif.

### *In vivo* bacterial abundance assays with *E. faecalis*

Mice were injected with either PBS (vehicle control) or 500 ng IL-25 every other day for seven days. On the day of the assay, mice were fasted for two hours and then given a single dose of 1×10^9^ *E. faecalis* strain OG1RF via oral gavage. Mice were euthanized and intestinal organs (small intestine, cecum, and large intestine) containing luminal contents were collected two hours post-gavage. Intestinal organs were individually homogenized in 2 mL of ice-cold PBS, serially-diluted in autoclaved MilliQ water, and cultured on MH-Rif agar overnight at 37°C to enumerate *E. faecalis* CFUs in each organ.

### Gut motility assays

Mice were injected with either PBS (vehicle control) or 500 ng IL-25 every other day for seven days. On the day of the assay, mice were fasted for two hours and then given a single dose of 1×10^6^ 10 μm Fluoresbrite® YG carboxylate microspheres (Polysciences) via oral gavage. At two hours post-gavage, mice were euthanized and luminal contents from the small intestine, cecum, and large intestine were collected. Microspheres were manually counted using a Nikon Eclipse E400 microscope and reported as total microspheres per organ.

### Statistical analysis

Statistical analysis of the spatial transcriptomics data and 16S rRNA sequencing data are reported in the corresponding Materials and Methods sections. Statistical tests used to compare differences between treatment groups in individual experiments are indicated in the figure legends. Student’s *t*-tests were used to determine statistical significance when a parametric test was appropriate; Mann-Whitney *U* tests were used when a non-parametric test was appropriate. Ordinary one-way ANOVA tests (and corresponding multiple comparison tests) were used to determine statistical significance in experiments that require comparisons of multiple experimental groups. A generalized linear model (GLM) with negative binomial distribution was performed to compare microbial count data. A linear mixed model (LMM) was used to determine statistical significance for quantitative western blot data. SAS 9.4 (SAS Institute Inc., Cary, NC) and GraphPad Prism were used to perform all statistical analyses. *p* < 0.05 was considered statistically significant. NS indicates no statistical significance, * *p* < 0.05, ** *p* < 0.01, *** *p* < 0.001, and **** *p* < 0.0001. GraphPad Prism software was used to generate all figures, unless otherwise indicated. Center values are arithmetic means, and error bars are standard error of the mean (SEM) in figures where Student’s *t*-tests, and ANOVA and LMM with multiple comparison adjustments were used to determine statistical significance. Center values are medians, and error bars are the interquartile range (IQR) in figures where Mann-Whitney *U* tests were used to determine statistical significance. For log-transformed data, center values are geometric means, and error bars are the 95% confidence interval (CI).

## Supplemental Figure Legends

**Figure S1.**
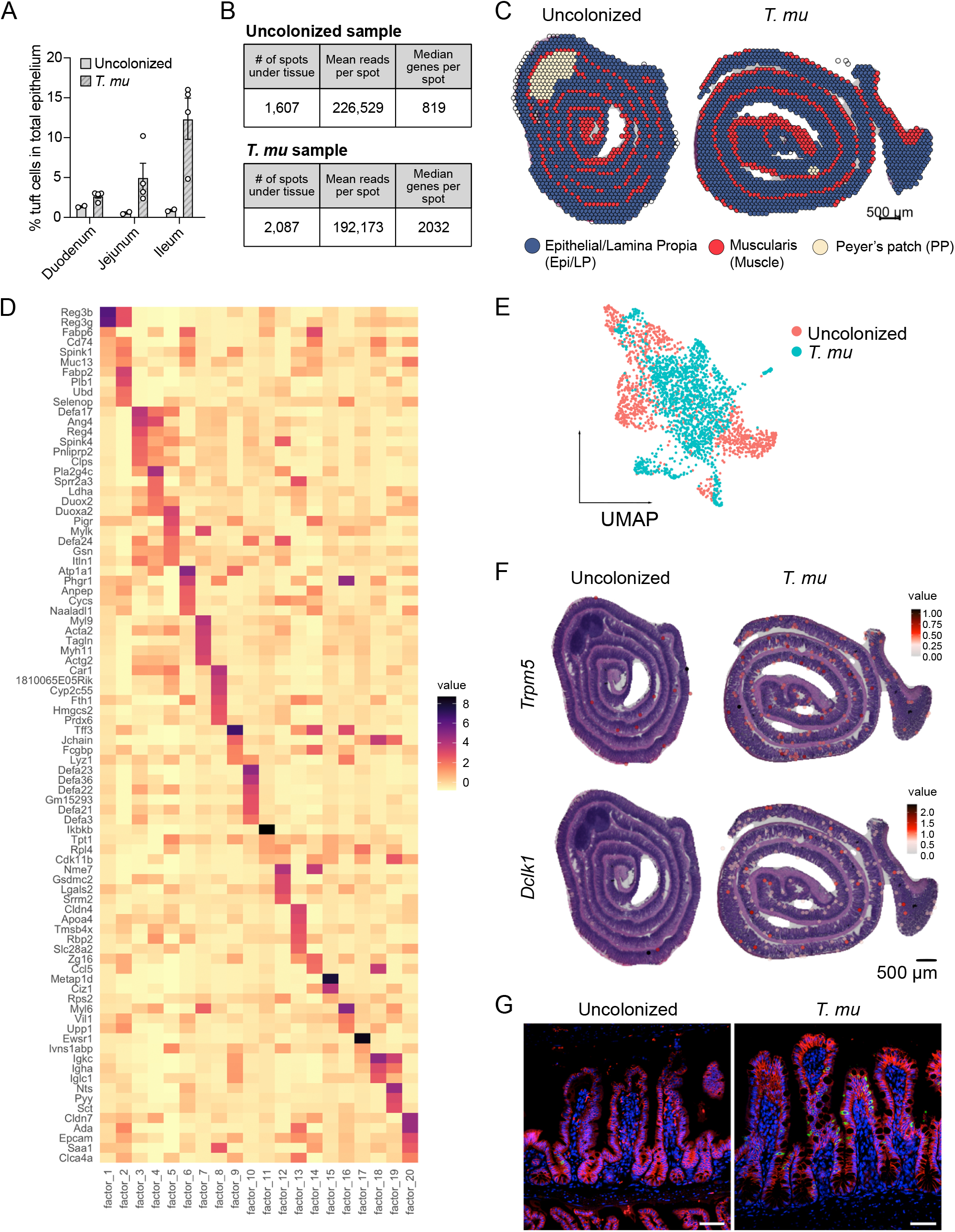
*T. mu* colonization induces profound small intestinal epithelial remodeling. (A) SI tuft cell frequency in uncolonized or *T. mu*-colonized *Gfi1b*^EGFP/+^ mice, in which tuft cells are specifically marked with EGFP, determined by flow cytometry (n = 2-4 mice per group). Center values = arithmetic mean; errors bars = SEM. (B) Summary of the number of spots sequenced, mean reads per spot, and the median genes per spot for both uncolonized and *T. mu*-colonized tissue sections that were used for spatial transcriptomics (ST) analysis. (C) Manual annotations of tissue compartments (Epithelial/Lamina Propria, Muscularis, or Peyer’s patch) overlaid onto the ST spots. (D) Non-negative matrix factor analysis on the uncolonized and *T. mu*-colonized ST samples. (E) Harmonized UMAP plot colored by sample identity to highlight the overlap of clusters shared between the treatment groups (uncolonized or *T. mu*-colonized). (F) Gene features overlay plots on ST spots throughout ileal tissue of the uncolonized or *T. mu*-colonized mouse, depicting expression of canonical tuft cell-specific genes, *Trpm5* and *Dclk1*. (G) Representative fluorescence microscopy images of the ileum from uncolonized or *T. mu*-colonized WT mice that were used for ST analysis. Nuclei (blue), E-cadherin (red), DCLK1 (green). Scale bar: 50 μm.

**Figure S2.**
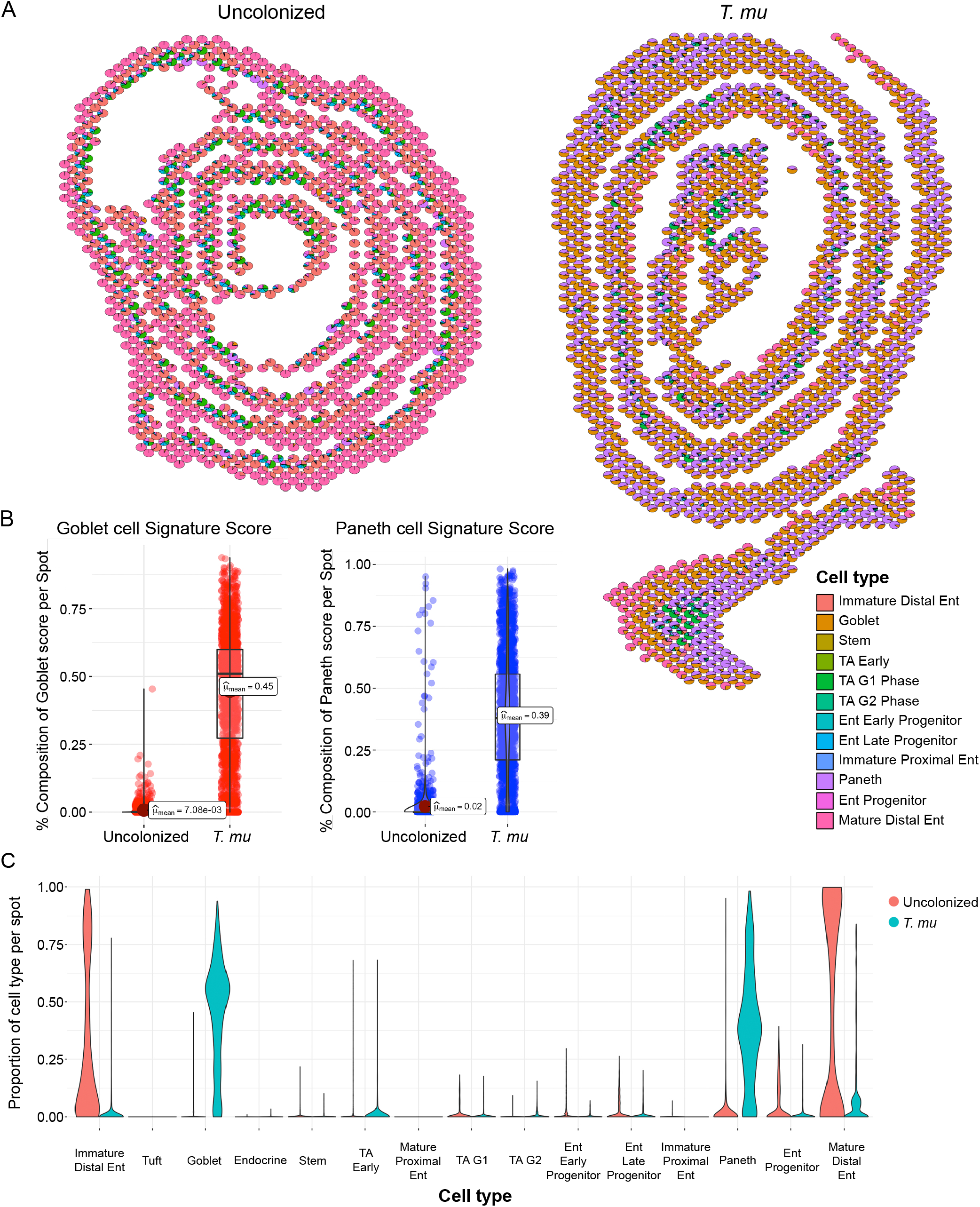
Single-cell RNA sequencing (scRNA-seq) integration reveals epithelial cell identity in ST spots. (A) Scatter pie plots of uncolonized (left) and *T. mu*-colonized (right) samples, colored by annotated cell identities. Ent, Enterocyte; TA, Transit-amplifying; G1, G1/S cell-cycle phase; G2, G2/M cell-cycle phase. (B) Box plot of ST spots scored on the composition of goblet (left; red) or Paneth cell (right; blue) score in the uncolonized and *T. mu*-colonized mice. (C) Violin plot of predicted epithelial cell types depicted for the two ST samples. Cell type identities were deconvoluted from scRNA-seq data from Haber et al., (2017) (32).

**Figure S3.**
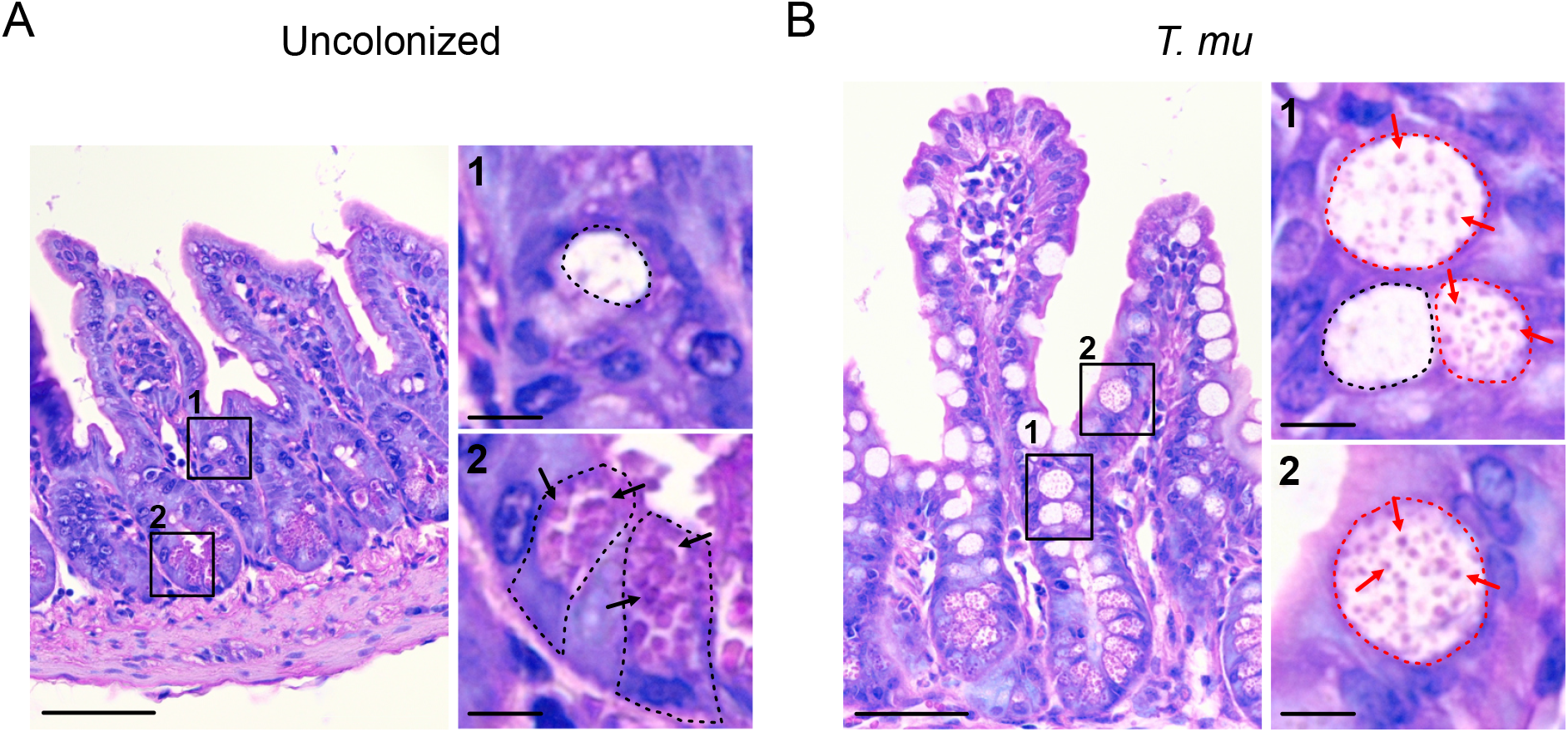
*T. mu* colonization increases the abundance of “intermediate cells” with hybrid goblet and Paneth cell morphologies. (A) Representative image of an H&E-stained section of several ileal crypt-villus axes from the uncolonized mouse used for ST analysis. Insets mark zoomed-in images to the right. 1: Dotted lines trace the outline of a goblet cell. 2: Dotted lines trace the outlines of individual Paneth cells. Arrows indicate representative Paneth cell secretory granules, which are round and stain light purple. Scale bar: 50 μm (original image), 10 μm (zoomed images). (B) Representative image of an H&E-stained section of several ileal crypt-villus axes from the *T. mu*-colonized mouse used for ST analysis. Insets mark zoomed-in images to the right. 1 and 2: Black dotted lines trace the outline of a goblet cell. Red dotted lines trace the outlines of individual “intermediate cells” with hybrid goblet-Paneth morphologies; red arrows indicate representative granules, which stain light purple, within these cells. Scale bar: 50 μm (original image), 10 μm (zoomed images).

**Figure S4.**
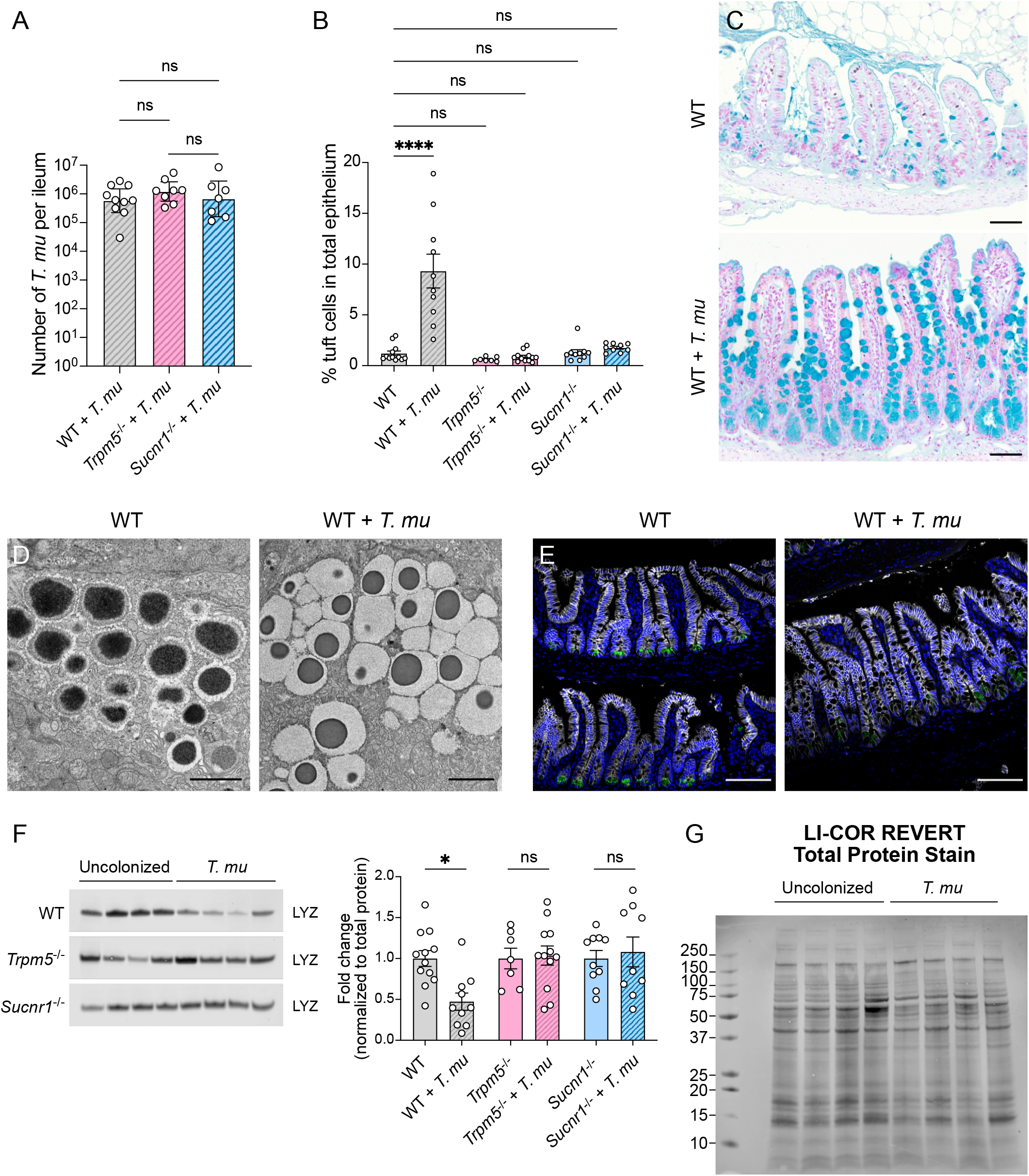
*T. mu* alters Paneth cell biology and AMP production by stimulating tuft cell taste-chemosensory pathways. (A) Quantitative PCR (qPCR) enumeration of *T. mu* abundance in the ilea of protist-colonized (3 weeks) WT, *Trpm5*^-/-^, and *Sucnr1*^-/-^ mice (n = 7-10 mice per group). Center values = geometric means; error bars = 95% CI. Significance determined using a generalized linear model (GLM). (B) Ileal tuft cell frequency in uncolonized or *T. mu*-colonized (3 weeks) WT, *Trpm5*^-/-^, and *Sucnr1*^-/-^ mice as determined by flow cytometry (n = 7-10 mice per group). Center values = geometric mean; error bars = 95% CI. Significance determined using an ordinary one-way ANOVA test. (C) Representative images of the ileum from uncolonized (top) or *T. mu*-colonized (3 weeks) WT mice (bottom), stained with Alcian blue and nuclear fast red to identify goblet cells. Alcian blue dye stains acidic polysaccharides; nuclear fast red stains nuclei. Scale bar: 50 μm. (D) Representative transmission electron microscopy images of secretory granules within Paneth cells in the ileal crypts from uncolonized or *T. mu*-colonized (3 weeks) WT mice. Scale bar: 2 μm. (E) Representative fluorescence microscopy images of uncolonized or *T. mu*-colonized (3 weeks) WT mouse ilea. Nuclei (blue), E-cadherin (white), Lysozyme (green). Scale bar: 100 μm. (F) Representative western blot images and quantitative analysis of intracellular lysozyme (LYZ) levels in the ilea of uncolonized or *T. mu*-colonized (3 weeks) WT, *Trpm5*^-/-^, and *Sucnr1*^-/-^ mice. Each band and symbol represents an individual mouse. LYZ levels were normalized to REVERT total protein stain (refer to Fig. S4G for an example image of REVERT staining corresponding to the blots) (n = 7-12 mice per group). Center values = arithmetic mean; error bars = SEM. A linear mixed model was used to determine significance. (G) Representative Western blot membrane of WT uncolonized or *T. mu*-colonized samples from Fig. S4F stained with REVERT total protein stain. REVERT total protein stain was used for normalization to determine relative intracellular LYZ levels in the different experimental samples. ns = no significance, * *p* < 0.05, **** *p* < 0.0001.

**Figure S5.**
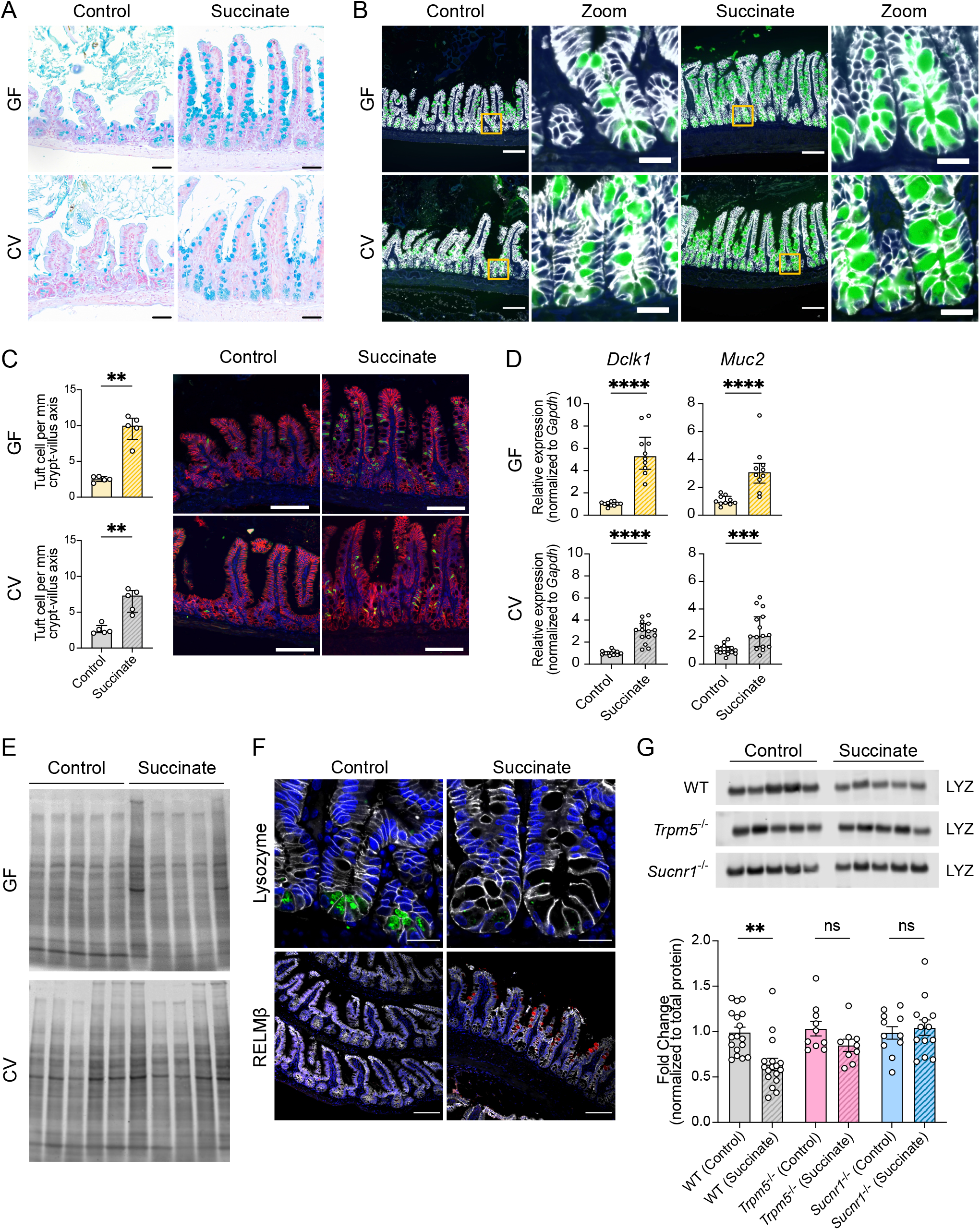
Succinate treatment led to tuft cell expansion and AMP changes in WT mice, but not in mice lacking tuft cell chemosensory components required to detect this metabolite. (A) Representative images of the ileum from control or succinate-treated (1 week) germ-free (GF) or conventionally colonized (CV) mice, stained with Alcian blue and nuclear fast red to identify goblet cells. Scale bar: 50 μm. (B) Representative fluorescence microscopy images of the ileum from control or succinate-treated GF or CV WT mice. Nuclei (blue), E-cadherin (white), MUC2 (green). Insets mark zoomed images (right). Scale bar: 100 μm (original image), 20 μm (zoomed images). (C) The number of tuft cells per mm of crypt-villus axis were quantified for control or succinate-treated (1 week) GF or CV mice (n = 5 mice per group). Center values are medians; error bars represent the interquartile range (IQR). Significance determined using Mann-Whitney *U* tests. Representative fluorescence microscopy images of each experimental condition are shown on the right. Nuclei (blue), E-cadherin (red), DCLK1 (green). Scale bar: 100 μm. (D) Expression of *Dclk1* (tuft cell marker) and *Muc2* (goblet cell marker) determined by RT-qPCR in the ilea of control or succinate-treated (1 week) GF and CV mice (n = 5-15 mice per group). Relative expression normalized to *Gadph*. Center values = median; error bars = IQR. Significance determined using Mann-Whitney *U* tests. (E) Representative western blot membranes of control or succinate-treated GF or CV WT mice (shown in Fig. 5D) stained with REVERT total protein stain. REVERT total protein stain was used for normalization to determine relative intracellular lysozyme levels in the different experimental samples. Images were cropped to remove samples that were unrelated to the experiment. (F) Representative fluorescence microscopy images of the ileum from control or succinate-treated (1 week) CV WT mice. Nuclei (blue), E-cadherin (white), Lysozyme (green), RELMβ (red). Scale bar: 20 μm (top images), 100 μm (bottom images). (G) Representative western blot images and quantitative analysis of intracellular lysozyme (LYZ) levels in the ilea of control or succinate-treated (1 week) WT, *Trpm5*^-/-^, and *Sucnr1*^-/-^ mice. Each band and symbol represents an individual mouse. LYZ levels were normalized to REVERT total protein stain (n = 9-17 mice per group). Center values = arithmetic mean; error bars = SEM. A linear mixed model was used to determine significance. ns = no significance, * *p* < 0.05, ** *p* < 0.01, *** *p* < 0.001, **** *p* < 0.0001.

**Figure S6.**
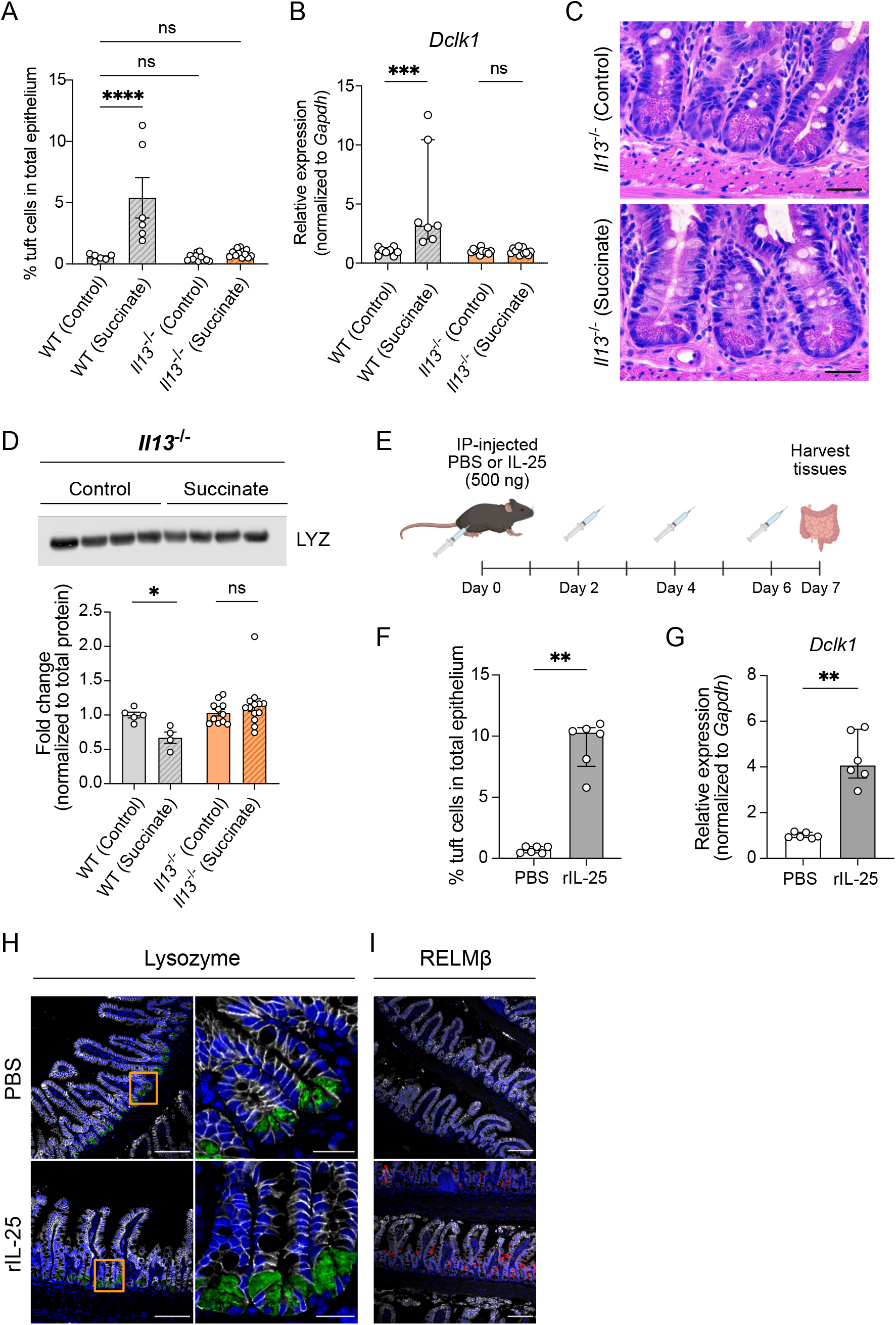
Type 2 immunity is required for changes to antimicrobial production in succinate-treated animals. (A) Ileal tuft cell frequency in control or succinate-treated (1 week) WT or *Il13*^-/-^ mice determined by flow cytometry (n = 6-12 mice per group). Center values = arithmetic mean; error bars = SEM. Significance determined using ordinary one-way ANOVA test. (B) Expression of *Dclk1* determined by RT-qPCR in the ileal epithelial fraction of control or succinate-treated (1 week) WT and *Il13*^-/-^ mice (n = 7-13 mice per group). (C) Representative H&E-stained images of the ileal crypts from control or succinate-treated (1 week) *Il13*^-/-^ mice. Scale bar: 25 μm. (D) Representative western blot images and quantitative analysis of intracellular lysozyme (LYZ) levels in the ilea of control or succinate-treated (1 week) WT and *Il13*^-/-^ mice. Each band and symbol represents an individual mouse. LYZ levels were normalized to REVERT total protein stain (n = 4-14 mice per group). Center values = arithmetic mean; error bars = SEM. A linear mixed model was used to determine significance. (E) Schematic of recombinant IL-25 injections. PBS (vehicle control) or 500 ng IL-25 were intraperitoneally (IP)-injected into mice every other day for one week, after which ileal tissues were harvested for downstream analyses. (F) Ileal tuft cell frequency in WT mice IP-injected with PBS or IL-25 determined by flow cytometry (n = 6 mice per group). Center values = median; error bars = IQR. Significance determined using Mann-Whitney *U* tests. (G) Expression of *Dclk1* determined by RT-qPCR in the ileal epithelial fraction of WT mice IP-injected with PBS or IL-25 (n = 6 mice per group). (H and I) Representative fluorescence microscopy images of the ileum from PBS or IL-25 injected WT mice. (H) Nuclei (blue), E-cadherin (white), Lysozyme (green). The insets mark zoomed in regions (right images). Scale bar: 100 μm (left images), 20 μm (right images). (I) Nuclei (blue), E-cadherin (white), RELMβ (red). Scale bar: 100 μm. ns = no significance, * *p* < 0.05, ** *p* < 0.01, *** *p* < 0.001, **** *p* < 0.0001.

**Figure S7.**
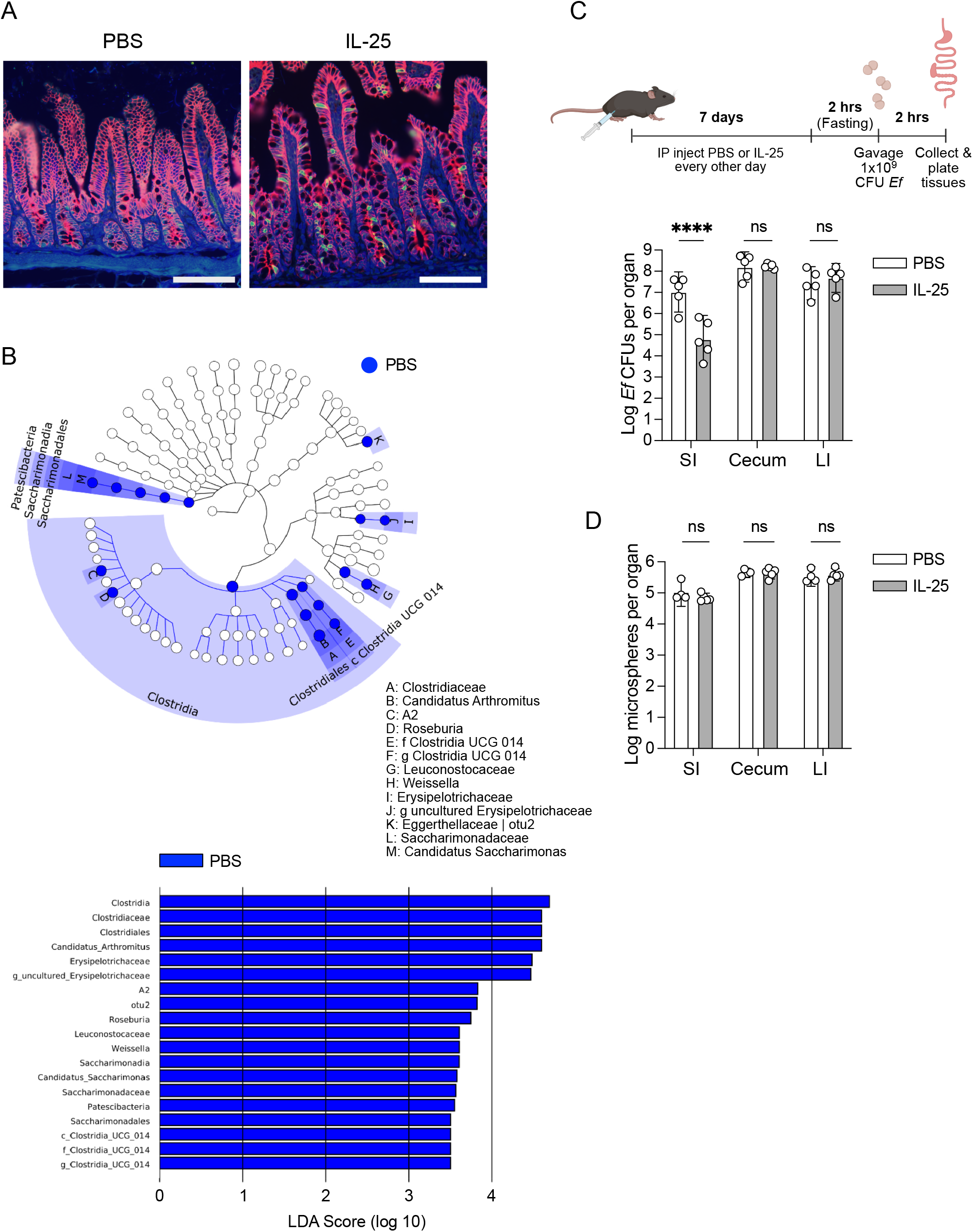
Recombinant IL-25 injections induce type 2 immunity and alter the ileal microbiome. (A) Representative fluorescence microscopy images of the ileum of PBS or IL-25 injected WT Taconic mice used for 16S rRNA sequencing experiment. Nuclei (blue), E-cadherin (red), DCLK1 (green). Scale bar: 100 μm. (B) Linear discriminant analysis (LDA) effect sizes (LEfSe) analysis using a LDA threshold score of 2 to identify ileal luminal bacterial taxa in PBS or IL-25 injected mice (n = 9-10 mice per group). Only PBS-injected mice showed differentially enriched taxa. Cladogram (top) highlights taxonomic relatedness of bacteria while LDA plot (bottom) is an ordered list of enriched bacteria. (C) Top: Experimental schematic. Mice were IP-injected with PBS or IL-25 as described in Fig. S6E, and then orally inoculated with *Enterococcus faecalis* (*Ef*). Bottom: *Ef* CFUs per organ were enumerated in the small intestine (SI), cecum, and large intestine (LI) (n = 5 mice per group). (D) Jackson mice were IP-injected with PBS or IL-25 as described in Fig. S6E, and then orally-inoculated with fluorescent microspheres. The luminal contents of the small intestine (SI), cecum, and large intestine (LI) were collected from each mouse at 2 hrs post-gavage for quantification of total microspheres per organ (n = 5 mice per group). In Panels C and D, center values = geometric mean; error bars = 95% confidence interval (CI). Significance determined using a generalized linear model. ns = no significance, **** *p* < 0.0001.

**Figure S8.**
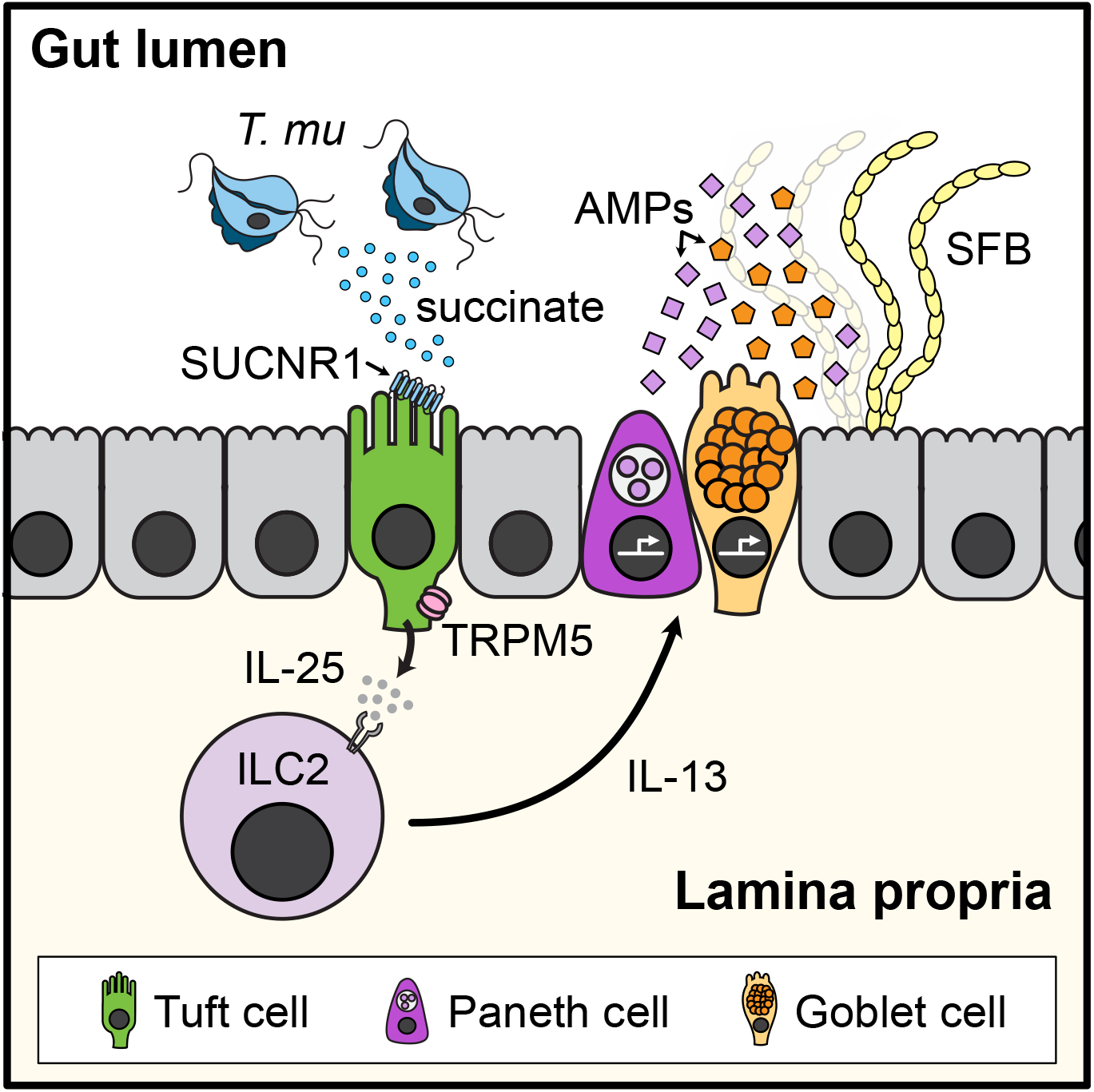
Model of how microbial-derived succinate remodels the small intestinal epithelial and antimicrobial landscape. Colonization with the commensal protist *T. mu* leads to tuft, goblet, and Paneth cell hyperplasia. This is due to *T. mu* excretion of succinate, which stimulates tuft cell taste-chemosensory signal transduction and eventually activates type 2 immunity. The type 2 cytokine IL-13 profoundly alters the AMP expression profile in the small intestine, leading to increased killing of mucosa-associated bacteria such as SFB.

**Figure S9.**
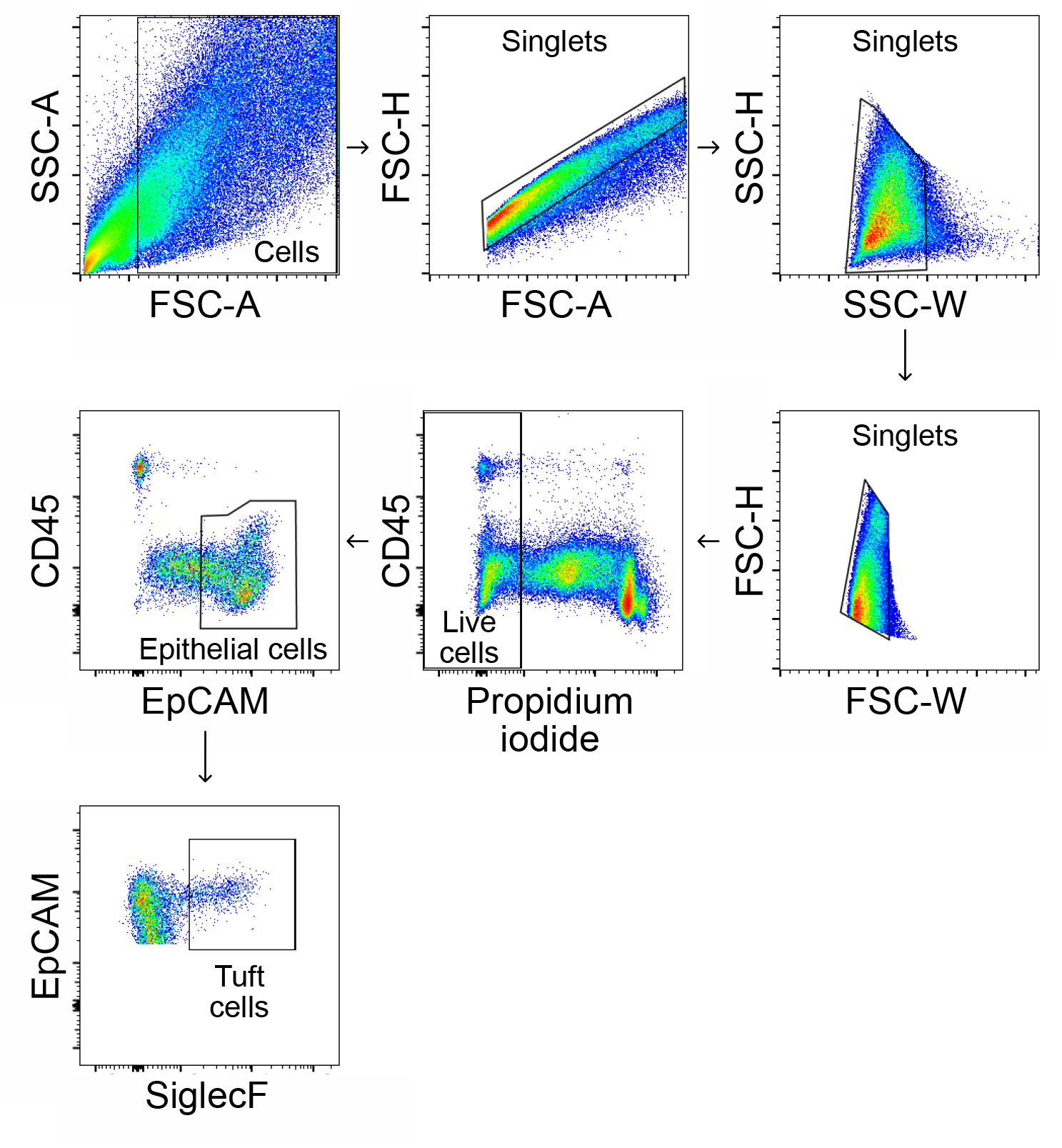
Tuft cell gating strategy for flow cytometry. Epithelial cells from the distal small intestine were isolated and gated on viable CD45^-^ EpCAM^+^ cells. Tuft cells were further selected as SiglecF^+^.

**Table S1.**
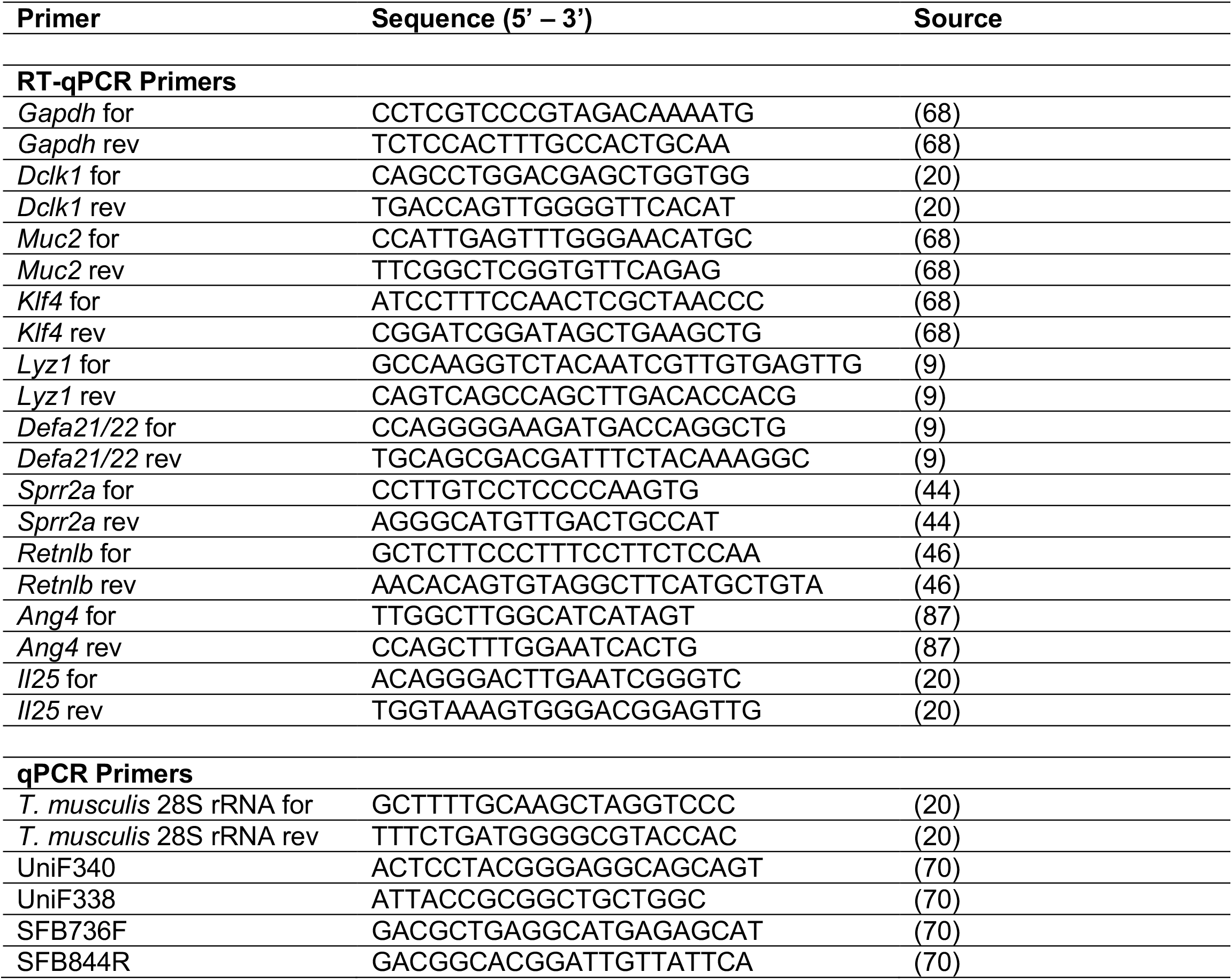
Primer sequences.

**Table S2.**
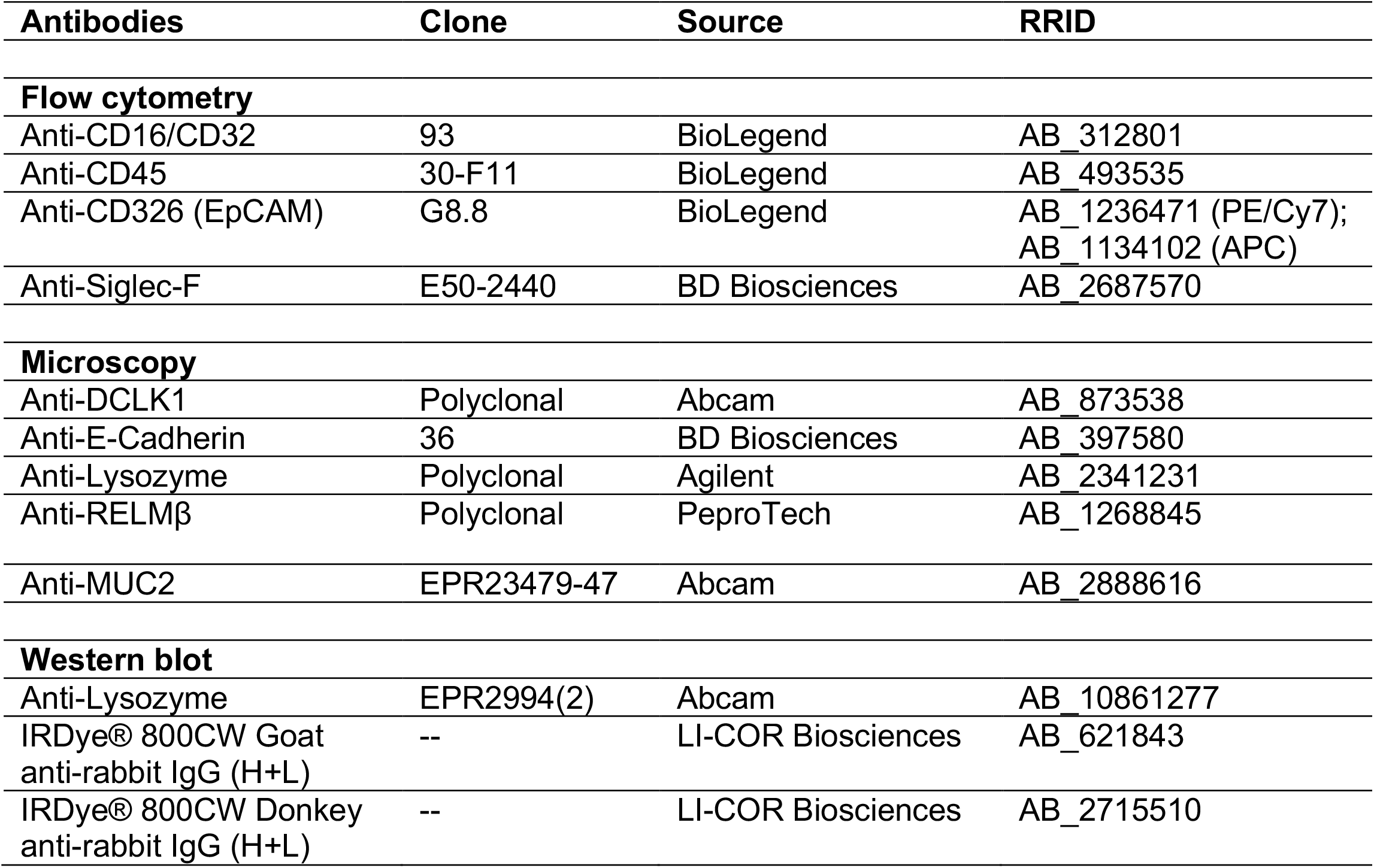
Antibodies and flow reagents.

